# EphA3 couples attractive guidance to persistent retinal axon growth through modulation of EphA4 forward signaling in a plasma membrane cholesterol-dependent manner

**DOI:** 10.64898/2026.09.15.751799

**Authors:** Gonzalo Spelzini, Mara Medori, Alejandro Scicolone, Sofia Martin Mena, Katia Del Rio-Tsonis, Luciano Fiore, Viviana Sanchez, Gabriel Scicolone

## Abstract

Topographic mapping requires retinal axons not only to avoid inappropriate targets but also to advance directionally through permissive and attractive territories. Using chick retinal ganglion cells in stripe and Dunn chamber gradient assays, we investigated how extracellular EphA3 coordinates the directional guidance and growth dynamics of nasal axons. Nasal axons preferentially grew on EphA3-containing stripes and turned toward increasing concentrations of EphA3 in soluble gradients. EphA3 also increased axonal growth velocity, primarily by prolonging extension phases and reducing growth interruptions rather than by increasing maximal extension-phase velocity. Pharmacological inhibition of EphA4 enhanced axonal extension but abolished EphA3-dependent attraction, indicating that basal EphA4 activity is required for spatial guidance even though EphA4 signaling can constrain axonal advance. Plasma membrane cholesterol depletion altered basal growth dynamics and attenuated EphA3-dependent turning and extension persistence. Together, these findings identify EphA3 as a noncanonical attractive cue that coordinates directional guidance with persistent nasal axon growth through mechanisms dependent on EphA4 activity and membrane cholesterol. They also suggest that sustained EphA4 inhibition may promote axonal growth while compromising topographic targeting.

**Highlights:**

- Environmental EphA3 attracts nasal RGC axons and promotes persistent forward advance.
- EphA3 prolongs extension phases rather than increasing maximal extension-phase velocity.
- Basal EphA4 activity is required for EphA3-dependent directional guidance.
- Plasma membrane cholesterol supports basal dynamics and the coordinated response to EphA3.
- Direct EphA4 inhibition dissociates axonal extension from positional guidance.

## INTRODUCTION

Topographic maps preserve spatial relationships between connected neuronal populations, and are essential for normal sensory function. In the retinotectal system, retinal ganglion cell (RGC) axons map the nasotemporal retinal axis onto the rostrocaudal axis of the optic tectum or superior colliculus (Fig. 1A). Successful mapping requires coordination between forward extension and selective responses to positional cues (Vanegas and Ito, 1983; Thanos and Bonhoeffer, 1987; Flanagan, 2006).

**Figure 1.**
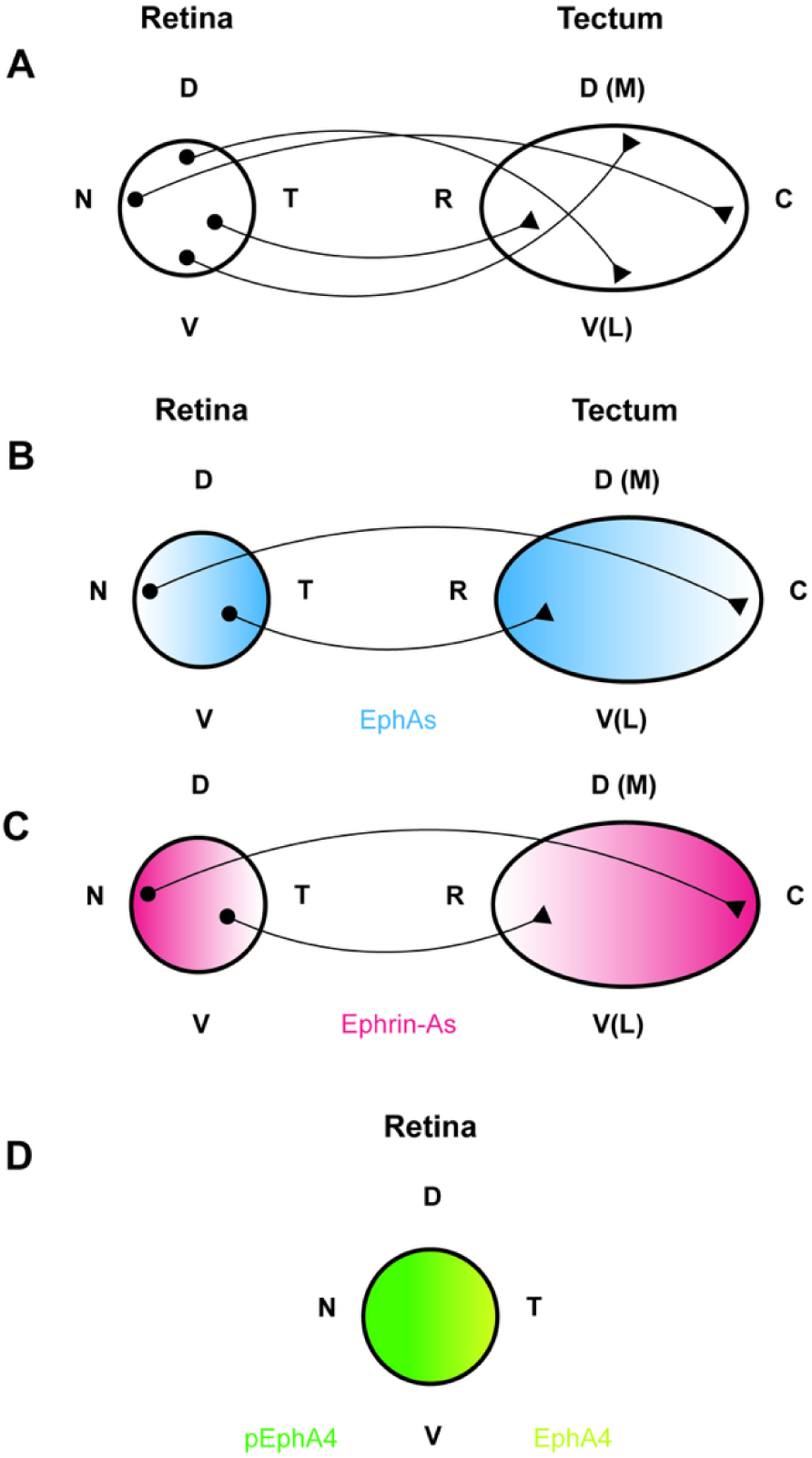
Schematic representation of retinotectal/retinocollicular topographic projections and EphA/ephrin-A expression gradients. (A) Topographic organization of retinal ganglion cell (RGC) projections along the nasotemporal and dorsoventral retinal axes. Nasal RGC axons project to the caudal tectum/superior colliculus (SC), whereas temporal RGC axons project to the rostral tectum/SC. Dorsal and ventral RGC axons project to the ventrolateral and dorsomedial tectal/collicular regions, respectively (N→C, T→R, D→V[L], and V→D[M]). Black dots represent RGC somata, and black arrowheads indicate their corresponding termination zones in the tectum/SC target regions. (B) EphA receptors are expressed at higher levels in the temporal retina and decrease toward the nasal retina. In the tectum/SC, EphA expression decreases along the rostrocaudal axis. (C) Ephrin-As are expressed at higher levels in the nasal retina and decrease toward the temporal retina, while their expression in the tectum/SC increases along the rostrocaudal axis. (D) Schematic representation of EphA4 distribution and activation along the nasotemporal retinal axis. EphA4 is expressed throughout this axis, whereas phosphorylated EphA4 (pEphA4) is higher toward the nasal retina (Ortalli et al., 2012). Color intensity represents relative expression or activation levels. Retinal region is indicated as nasal (N), temporal (T), dorsal (D), and ventral (V); tectal/collicular area is indicated as rostral (R), caudal (C), dorsomedial (D[M]), and ventrolateral (V[L]). Modified from Scicolone et al., 2009.

Canonical models emphasize the repulsive effect of the increasing caudal gradient of tectal ephrin-As on temporal RGC axons, which express higher levels of EphA receptors. This repulsion excludes temporal axons from caudal tectal territories and favors their branching and termination near their rostral targets (Cheng et al., 1995; Drescher et al., 1995; Feldheim et al., 2000; Monschau et al., 1997; Nakamoto et al., 1996; Sakurai et al., 2002; Yates et al., 2001). Repulsive exclusion alone, however, does not fully explain how nasal axons are actively guided into and across the tectum while maintaining their growth and positional selectivity ( Hornberger et al., 1999; Scicolone et al., 2009; Medori et al., 2020).

Previous work has shown that tectal EphA3 can bind axonal ephrin-As, reduce ephrin-A-dependent EphA4 forward signaling, stimulate nasal RGC axon growth, and restrict premature branching in rostral tectum (Ortalli et al., 2012; Fiore et al., 2019; Medori et al., 2020). These observations raise the possibility that substrate-bound EphA3 acts noncanonically as an attractive ligand. However, the real-time growth cone dynamics underlying this response and the contributions of basal EphA4 activity to EphA3-mediated axon growth and guidance remain incompletely understood. Moreover, reduced EphA4 expression has been associated with increased growth of partially regenerated axons, suggesting that EphA4 signaling may also be relevant to axonal regeneration (Chen et al., 2022; Ning et al., 2020).

Ephrin-As are constitutively located in lipid rafts, whereas EphA receptors are recruited to these domains following ephrin-A binding (Davy et al., 1999; Gauthier et al., 2003; Averaimo et al., 2016). Previous studies have examined the influence of cholesterol-rich membrane microenvironments on axon growth and guidance (Ibáñez, 2004; Guirland et al., 2004; Viljetić et al., 2024). However, the role of plasma membrane cholesterol in EphA3-mediated axon growth and guidance remains unknown. Furthermore, transient cholesterol depletion with β-methylcyclodextrin (βMCD) has produced contrasting effects across different experimental models (Guirland et al., 2004; Zidovetzki and Levitan, 2007; Griswold et al., 2025).

We hypothesized that tectal EphA3 acts as an attractive guidance cue by modulating axonal EphA4 signaling within cholesterol-dependent membrane domains, thereby coupling directional turning to persistent forward growth. To test this hypothesis, we combined EphA3 substrate-choice assays with Dunn chamber gradients of EphA3 and time-lapse analyses of axonal turning and growth dynamics, including net advance, extension, pausing, and retraction, to evaluate nasal RGC axon behavior. We also inhibited EphA4 signaling pharmacologically with KYL (Fiore et al., 2019; Lamberto et al., 2012; Murai et al., 2003; Noberini et al., 2012) and depleted plasma membrane cholesterol with βMCD to assess the roles of EphA4 activity and membrane cholesterol integrity in nasal RGC axon growth and guidance.

## MATERIALS AND METHODS

### Animals

Pathogen-free fertilized White Leghorn chicken (Gallus gallus) eggs were obtained from Rosenbusch Institute (Buenos Aires). They were incubated at 38 °C and 60% relative humidity. Developmental stages were recorded according to Hamburger and Hamilton Normal Table of Stages (HH) (Hamburger and Hamilton, 1951). Animals were treated following the Guide for the Care and Use of Laboratory Animals from the Institute of Laboratory Animals Resources, Commission of Life Sciences, National Research Council, USA, and approved by the Council for Care and Use of Experimental Animals from University of Buenos Aires.

### Cultures of dissociated retinal neurons and retinal explants

Seven days embryos (E7) (HH 30–31) were removed from the eggs, decapitated and their retinas were dissected out in ice-cold HBSS (pH 7.4). These stages of development were chosen since dissociated retinal cultures and explants obtained from these ones, present the best axonal growth in vitro (Cohen et al., 1989; Halfter et al., 1981) and coincide with the period in which RGC axons invade the tectum (Feldheim and O’Leary, 2010; Scicolone et al., 2009).

The cornea, lens, marginal zone and vitreous humor were removed and discarded, exposing the neural retina which was separated from the retina pigmented epithelium. Retinas were divided into three sections from nasal to temporal pole, middle third was discarded and nasal or temporal thirds were employed.

For cultures of dissociated retinal neurons, the neural retinal portions were mechanically and enzymatically dissociated with a solution containing trypsin (0.1%), EDTA (1mM) and DNAse (0.05%). The cells were resuspended in N2-supplemented (Invitrogen) F12/DMEM medium (Invitrogen) and cultured at 10,000/15,000 cells/cm2 on coated coverslips located onto culture dishes (Ortalli et al., 2012).

Retinal explants were obtained by cutting small pieces from nasal or temporal retinas, and cultured in N2-supplemented F12/DMEM containing 0.4% methylcellulose (Sigma) on coated coverslips located onto culture dishes (Ortalli et al., 2012). Substrates were prepared by incubating coverslips with poly-L-lysine (100 μg/ml) (Sigma) and laminin (20 μg/ml) (Invitrogen) for 2 h at room temperature (RT) each one.

Both the dissociated retinal neurons and the retinal explants were cultured at 37 °C and 5% CO2 for 24 h. Afterward, the cultures were fixed with paraformaldehyde 2%-sacarose 2% in PBS for 30 min at RT (Ortalli et al., 2012).

### Stripe Assay

To analyze axonal responses to surface-bound guidance cues, a modified protein stripe assay was performed based on previously described protocols. Alternating protein stripes were generated on poly-L-lysine-coated culture dishes using silicone microchannel matrices. Briefly, an Fc-containing protein solution, combined with fluorescently labeled anti-Fc antibody, was first applied to the matrix to generate the stamped stripe pattern. After incubation and washing, the matrix was transferred onto the culture surface, and a second Fc-tagged protein solution was applied through the microchannels, allowing the formation of alternating, physically separated protein stripes (Gebhardt et al., 2012; Weschenfelder et al., 2013).

EphA3-Fc or Fc control proteins were used at 0.25 μg/ml and clustered with anti-Fc antibody at 0.085 μg/ml. Fluorescently labeled anti-Fc antibody was used to visualize the stripe pattern. After removal of the matrix, the patterned surface was washed and coated with laminin at 20 μg/ml to promote axonal growth. Stripe quality and continuity were verified by fluorescence microscopy before use.

Retinal explants were plated onto the patterned substrates and cultured under the corresponding experimental conditions. When indicated, cultures were treated with the EphA4 inhibitor KYL peptide at a final concentration of 13 μM (Lamberto et al., 2012; Murai et al., 2003; Noberini et al., 2008, 2012). Control cultures received an equivalent volume of culture medium. Axonal growth and growth cone morphology were analyzed after incubation.

### Morphometric analysis

After 24 h of incubation, retinal explants were imaged by differential interference contrast (DIC) microscopy and epifluorescence using 10× and 20× objectives. Images of the explants, axonal outgrowth, and fluorescent stripe patterns were stitched and manually aligned using Adobe Photoshop (Adobe Inc., San Jose, CA, USA; various versions used between 2019 and 2024).

Axonal preference was analyzed on merged DIC/fluorescence images using FIJI/ImageJ (Schindelin et al., 2012).

To quantify axonal distribution on the patterned substrates, individual axons were manually classified according to their final position and trajectory relative to the alternating stripes. Regions of interest were used to trace axons and assign them to one of three categories: axons ending on the chemoattractant stripe, axons ending on the non-attractant stripe, or axons showing no clear preference by crossing two or more different stripes. For each explant, the number of axons in each category was quantified and expressed as a percentage of the total number of analyzed axons.

Explants were then assigned a final preference score based on the predominant axonal category: 1, preference for the chemoattractant stripe; -1, preference for the non-attractant stripe; or 0, no evident preference. This individual axon-based classification was used to provide a more objective quantification of axonal preference compared with global qualitative scoring of the entire explant (Walter et al., 1987; Feldheim et al., 2000).

The axonal preference was also evaluated by using a preference score based on colour classification. Thus, white axons (without preference) were assigned a value of 0 (cero), green axons (preference for labeled anti-FC lanes) were assigned a value of -1 (less one) and yellow axons (preference for EphA3-Fc or unlabeled anti-Fc) were assigned a value of +1 (plus one). (Fig. 2 A-D, Supplementary Fig. 1 A-D).

**Figure 2.**
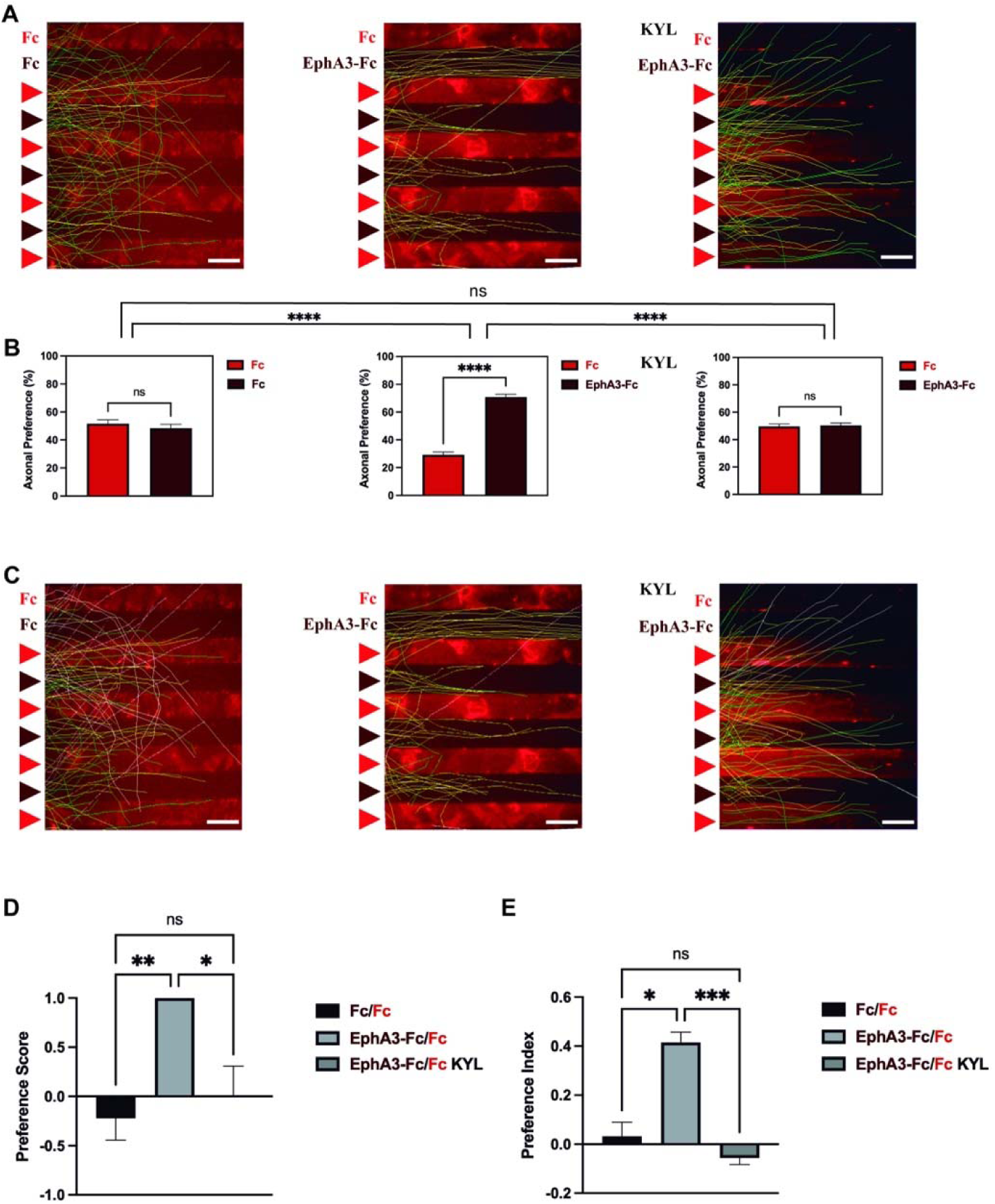
Nasal RGC axons preferentially grow on clustered EphA3-Fc stripes and this preference depends on modulation of axonal EphA4 forward signaling. A and C. Merged images of phase contrast and epifluorescent microphotographs of nasal RGC axons grown from explants obtained from E7 (HH30-31) chicken nasal retina cultured in three different stripe assays are shown: Left: anti-Fc (obscure) vs labeled anti-Fc (Alexa 594, red); Middle and right; EphA3-Fc clustered with anti-Fc (obscure) vs labeled anti-Fc (Alexa 594, red) without (middle) or with KYL (right). Axons were manually reconstructed to evaluate their distributions. In (A) the axons were classified according to their final position (yellow: preference for EphA3-Fc or unlabeled anti-Fc corridors, green: preference for labeled anti-Fc lanes). In (C) the axons were subclassified according to their final position and their trajectories (white: without preference because they cross through more than one limit between lanes, yellow: preference for EphA3-Fc or unlabeled anti-Fc corridors, green: preference for labeled anti-Fc lanes). Bar = 50 μm. B. Bar graphs represent the percentages of axons whose growth cones are located on anti-Fc labeled with Alexa 594 vs unlabeled anti-Fc stripes (left), the percentages of axons that are located on the anti-Fc labeled with Alexa 594 vs EphA3-Fc clustered with anti-Fc (middle), and the percentages of axons that are located on the anti-Fc labeled with Alexa 594 vs EphA3-Fc clustered with anti-Fc with presence of KYL (right). Data are shown as mean ± SEM. Significance of difference between lanes inside each condition was assayed by Welch t-test (for normal distribution) or Mann–Whitney test (for a non-normal distribution). Comparisons among different experimental conditions were assayed by Kruskal– Wallis and Dunn posttest. (D) Bar graph represents the axonal preference score which was calculated for each explant (0 = without preference, -1 = preference for labeled anti-Fc, 1 = preference for EphA3-Fc or anti-Fc). (E) Bar graph represents the axonal preference index calculated as (*N* EphA3-Fc or anti-Fc − *N* labeled anti-Fc)/(*N* EphA3-Fc or anti-Fc + *N* labeled anti-Fc). Data in (D) and (E) are shown as mean ± SEM. Significance of differences among different experimental conditions was assayed by Kruskal– Wallis and Dunn posttest (*p < 0.05; **p < 0.01; ***p < 0.001; ****p < 0.0001; ns, not significant). These are the same for all figures. Minimum N of axons for condition: 50, minimum N of explants for condition: 7.

Also, a preference index was calculated (N EphA3-Fc or anti-Fc − N labeled anti-Fc)/(N EphA3-Fc or anti-Fc + N labeled anti-Fc), which normalizes the magnitude of preference into a range between -1 to 1. (Fig. 2 E).

### Dunn’s chamber chemotaxis assay

Chemotaxis assays were performed using Dunn’s chambers, adapted from previously described protocols (Wells et al., 2005; Yam et al., 2009; Benseddik et al., 2022). Dissociated retinal cells from E7 chick embryos were plated on poly-L-lysine/laminin-coated coverslips and cultured for 12 h before the assay. Coverslips were then inverted onto the Dunn’s chamber, allowing cells located over the annular bridge between the inner and outer wells to be imaged.

The inner well was filled with a complete culture medium consisting of F12/DMEM supplemented with 1% N2. The outer well was filled with a complete medium containing the corresponding guidance cue, thereby generating a stable gradient across the bridge by diffusion.

The kinetics of gradient formation in the Dunn chamber are predictable based on diffusion and have been validated using fluorescent dyes (Zicha et al., 1991).

Accordingly, the molecular weight of each guidance molecule was taken into account when determining the gradient-establishment period before time-lapse imaging was initiated.

The following conditions were used: 1) EphA3-Fc/anti-Fc gradient -preclustered for 1 h at 37°C at a 2:1 molar ratio, with EphA3-Fc used at 0.25 μg/ml; 2) βMCD gradient at 5 mM (Zidovetzki & Levitan, 2007); 3) EphA3-Fc/anti-Fc gradient with βMCD; 4) EphA3-Fc/anti-Fc with KYL peptide at a final concentration of 13 μM (Lamberto et al., 2012; Murai et al., 2003; Noberini et al., 2008, 2012); and 5) control condition (complete medium alone was added to the outer well).

After chamber assembly, time-lapse imaging was performed at 37°C. Following 2 h of incubation, cultures were fixed with 2% paraformaldehyde in 2% sucrose to preserve growth cone morphology. Retinal ganglion cells were subsequently identified by post-hoc Islet-2 immunostaining, allowing the analyzed neurons to be reassigned to their corresponding experimental condition and cellular identity.

### Time-lapse microscopy

Time-lapse imaging was performed immediately after Dunn’s chamber assembly. The environmental chamber was pre-equilibrated at 37°C before imaging, and the Dunn’s chamber was allowed to thermally equilibrate on the microscope stage for 15 min. Cells located over the annular bridge region were imaged, and fields were selected to include the outer edge of the bridge in order to determine the direction of the chemical gradient.

Phase-contrast images were acquired using an Olympus IX83 inverted microscope equipped with a 10× objective. For each experiment, 5-6 fields were recorded for 2 h at 4-min intervals, using a 50-ms exposure time. Illumination was kept off between acquisitions to minimize phototoxicity. Image sequences from each field were saved as individual time-lapse stacks for subsequent analysis.

### Data analysis

Time-lapse image sequences were analyzed to quantify axonal growth, turning behavior and growth cone dynamics. Image stacks were generated and analyzed using FIJI/ImageJ. Axons showing net growth during the imaging period were selected, and individual regions of interest were manually traced from the soma to the growth cone at the first and last frames of the recording. Axonal length at 0 and 120 min was measured after spatial calibration, and net axonal growth was calculated as the difference between final and initial length.

Axonal guidance was analyzed using GeoGebra Classic (GeoGebra, 2025). For each selected axon, the position of the field within the Dunn’s chamber bridge was reconstructed to determine the direction of the gradient. The initial angle α was defined as the angle between the initial axonal orientation and the direction of the gradient. The turning angle β was defined as the angle between the initial axonal position and the displacement vector of the growth cone from the first to the last frame. Positive values indicated turning toward the gradient, whereas negative values indicated turning away from it.

Only non-fasciculated axons with an initial angle α between 0° and 180° and a net extension of at least 10 μm during the recording period were included in the analysis. Axons were excluded if they contacted debris, other cells, neighboring axons or their own soma. The initial position of the growth cone was defined at time 0.

Growth cone trajectories and axonal dynamics were further analyzed in FIJI using the Manual Tracking plugin. Growth cone position was manually tracked throughout the 2 h recordings at 4-min intervals. For each axon, XY coordinates, frame-to-frame displacement, total distance travelled and average velocity were obtained. Forward growth was assigned positive values, retraction was assigned negative values, and pauses were scored as zero displacement. This analysis allowed the quantification of axonal growth, turning and velocity in relation to the position of each axon within the chemotactic gradient.

### Immunocytochemistry

Dissociated retinal cells were fixed with 2% paraformaldehyde and 2% sucrose in PBS for 30 min at room temperature (RT) and rinsed with PBS. Non-specific binding was blocked by incubation with 5% normal goat serum (NGS) in PBS, with 0.5% Tween 20, for 1 h. Cultures were then incubated overnight at 4°C with mouse monoclonal antibodies against Islet-2 (clone 51.4H9, mouse IgG1; Developmental Studies Hybridoma Bank, DSHB; RRID:AB_528316), used as a retinal ganglion cell marker, or βIII-tubulin (clone 6G7, mouse IgG1; DSHB; RRID:AB_528497), used as a neuronal marker. Both primary antibodies were diluted to a final concentration of 2 μg/ml in PBS containing 1% NGS.

After washing with PBS, cultures were incubated for 2 h at RT, protected from light, with Alexa Fluor 488- or Alexa Fluor 594-conjugated F(ab′) □ fragments of goat anti-mouse IgG antibodies (A-11017 and A-11020, respectively; Molecular Probes), diluted 1:1000 (1 μg/ml) in PBS containing 1% NGS. Nuclei were counterstained with Hoechst 33342 diluted 1:10,000 for 10 min. Negative controls were processed in parallel by omission of the primary antibody.

Cultures were subsequently rinsed with PBS and mounted with Fluoromount-G (SouthernBiotech). Images were acquired using an Olympus IX83 inverted microscope. When indicated, the actin cytoskeleton of RGC axons was labeled with Alexa Fluor 488-conjugated phalloidin (A-12379, Molecular Probes).

Immunofluorescence for Islet-2 and βIII-tubulin, together with Hoechst nuclear staining, was used to characterize dissociated retinal cells and identify retinal ganglion cells and neuronal populations.

### Statistical analysis

All the quantifications were made in blind to experimental conditions. Data were expressed as mean ± SE. ANOVA and Tukey posttest or Student’s t-test were used for comparisons of axon length and intensity of labeling in immunocytochemistry when samples presented normal distributions. Non parametric Kruskal-Wallis and Dunn’s posttest or Mann Whitney test were employed when samples did not present normal distributions. When values were normalized to control conditions, ANOVA and Dunnett posttest were employed. p < 0.05 was considered significant. The cumulative distributions of turning β angles for each condition were compared statistically using the two-sample Kolmogorov-Smirnov test. p < 0.05 was considered significant. The Python notebooks used for the statistical analyses and figure-generation workflows described in this study are publicly available at https://github.com/gspelzini/axonaldynamicsandguidance. A versioned snapshot of the repository (v1.0.0) has been archived in Zenodo and is available at https://doi.org/10.5281/zenodo.22711392. The repository includes synthetic example datasets solely to demonstrate and test the workflows. These synthetic datasets do not contain experimental observations and are not intended to reproduce the numerical results reported in this study.

## RESULTS

### 1. Substrate-bound EphA3 attracts nasal RGC axons through modulation of EphA4 signaling

Nasal RGC axons were tested on alternating stripes of clustered EphA3-Fc and fluorescent anti-Fc; whereas alternating labeled and unlabeled anti-Fc stripes served as controls. Because EphA3-dependent growth stimulation has been linked to reduced ephrin-A-mediated EphA4 forward signaling, parallel cultures were treated with the EphA4 antagonist KYL (Fiore et al., 2019; Lamberto et al., 2012; Murai et al., 2003; Noberini et al., 2012).

Nasal RGC axons preferentially grew on clustered EphA3-Fc stripes but showed no preference between control stripes (Fig. 2A-C). KYL abolished EphA3-Fc stripe preference, while control stripes remained neutral (Fig. 2A-C; Supplementary Fig. 1).

Both quantitative measures confirmed this response. Preference scores approached +1 on EphA3-Fc stripes but remained near 0 under the control and EphA3-Fc + KYL conditions (Fig. 2D). Likewise, the preference index approached 0.4 with EphA3-Fc and 0 under the other conditions (Fig. 2E). Thus, substrate-bound EphA3 attracts nasal RGC axons, whereas direct EphA4 inhibition eliminates rather than reproduces this preference, indicating that attraction requires a regulated level of EphA4 signaling.

### 2. EphA3 attracts nasal RGC axons through mechanisms requiring plasma membrane cholesterol and basal EphA4 activity

Stripe assays test choices between fixed substrate concentrations, whereas RGC axons encounter molecular gradients *in vivo*. To better mimic *in vivo* conditions, we used Dunn chambers and time-lapse imaging to determine whether clustered EphA3-Fc gradients attract nasal RGC axons and whether this response requires membrane cholesterol or EphA4 activity. Dissociated E7 chick retinal neurons were imaged every 4 min for 120 min. Neurons were recognized throughout their morphology and by immunolabeling against neuron specific bIII tubulin. Only the longest neurite of each neuron exceeding twice the cell body diameter was considered as a RGC axon and taken into account for quantification (Ortalli et al., 2012). RGC identity was confirmed by Islet-2 immunolabeling (Supplementary Fig. 2). The turning angle β was defined as positive toward the gradient and negative away from it.

Analysis of the mean angular deviation β (Fig. 3A-C,G) showed that the clustered EphA3-Fc gradient induced attractive turning of nasal RGC axons (≈ +20 °), and differed significantly from the non-gradient control conditions (≈ 0°; p < 0.05). Representative time-lapse sequences showed growth-cone trajectories toward the source of EphA3 gradient (Fig. 3G). Cholesterol depletion with βMCD in the absence of a molecular gradient did not significantly alter turning relative to the control (Δ ≈ +3 °, ns). βMCD treatment attenuated EphA3-Fc-induced attractive turning ( ≈ +10 °), producing an intermediate response that did not differ significantly from either the controls or the EphA3-Fc gradient alone (Fig. 3C). Inhibition of ephrin-A-mediated EphA4 forward signaling with KYL abolished EphA3-Fc-induced attractive turning ( ≈ 0 °) (Fig. 3E,G).

**Figure 3.**
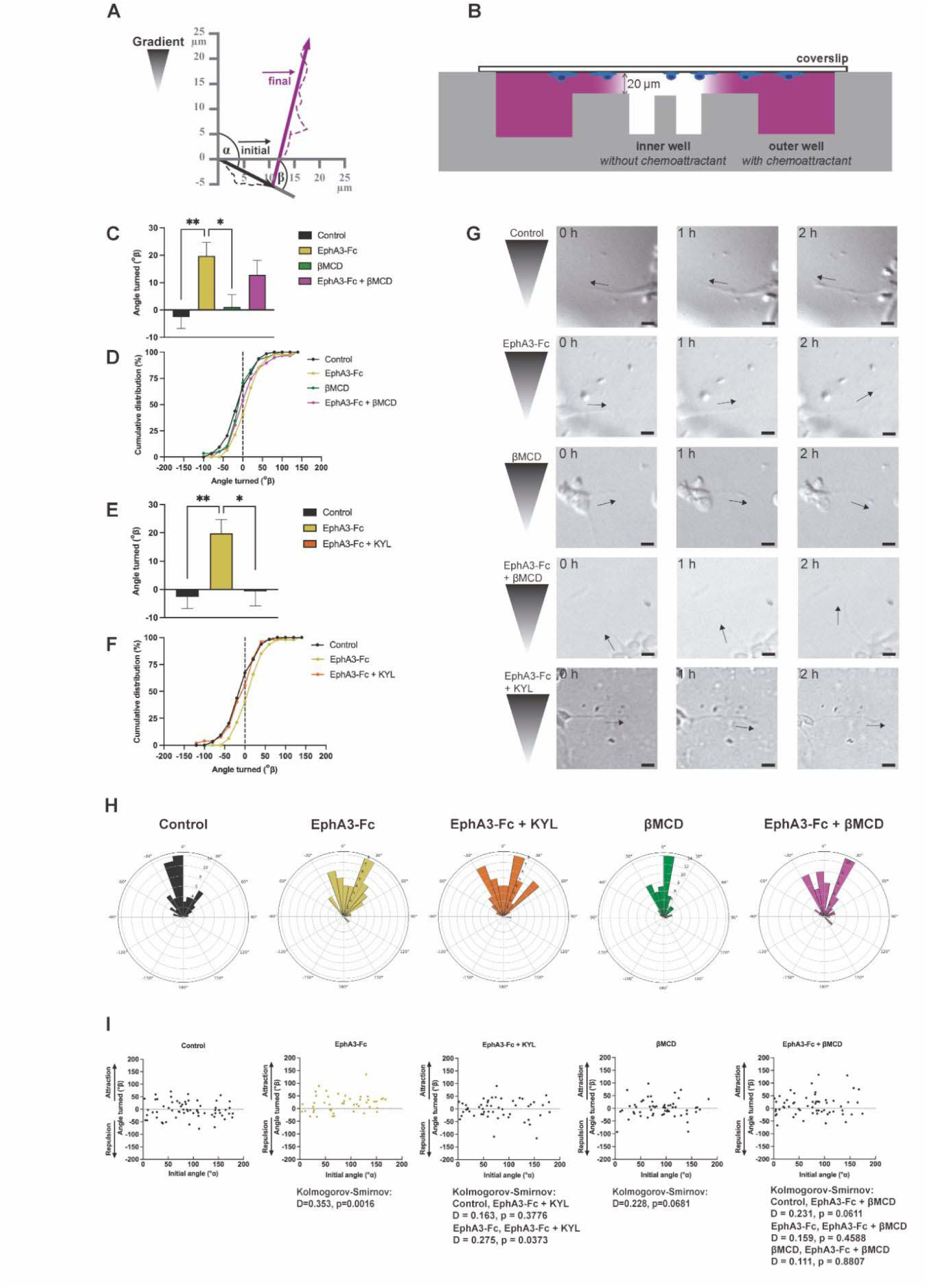
The chemoattractant effect of a clustered EphA3-Fc gradient on nasal RGC axons is abrogated by inhibition of axonal EphA4 signaling and partially inhibited by cholesterol depletion with βMCD. (A) Schematic illustrating how the initial and final directions of axon extension were used to calculate the angle turned (β) relative to the chemoattractant gradient. (B) Schematic cross-section of the Dunn chemotaxis chamber, showing the inner and outer wells, the coverslip, and the chemoattractant gradient generated across the bridge between the wells. (C, E) Mean angle turned (β°) by growth cones under the indicated experimental conditions. A positive β value indicates turning toward the gradient (attraction), whereas a negative value indicates turning away from it (repulsion). Bars represent mean ± SEM. Multiple comparisons are indicated above the bars; Kruskal–Wallis followed by Dunn’s test: *p < 0.05, **p < 0.01; ns, not significant. (D, F) Cumulative distributions of the turning angles under the corresponding conditions; the dashed vertical line at 0° indicates no net attraction or repulsion. (G) Representative DIC time-lapse micrographs at 0, 1, and 2 h of axons in a Dunn chemotaxis chamber under each condition. A schematic of the applied gradient direction is shown to the left of each row, and the black arrow indicates the main axis of the axon analyzed. Scale bar: 10 μm. (H) Polar histograms showing the angular distribution of growth-cone turning under each condition, grouped according to the sign of the angle; 0° represents no turning, positive angles indicate attraction, and negative angles indicate repulsion relative to the gradient. (I) Scatter plots relating the initial growth-cone angle (x-axis) to the angle turned (y-axis) for each condition; the horizontal dotted line indicates 0° (no deflection). Each panel reports the Kolmogorov–Smirnov test values (D, p) comparing the β distributions of the indicated conditions: Control versus EphA3-Fc (D = 0.353, p = 0.0016), βMCD (D = 0.228, p = 0.0681), EphA3-Fc + βMCD (D = 0.231, p = 0.0611), and EphA3-Fc + KYL (D = 0.163, p = 0.3776); EphA3-Fc versus EphA3-Fc + βMCD (D = 0.159, p = 0.4588) and EphA3-Fc + KYL (D = 0.275, p = 0.0373); and βMCD versus EphA3-Fc + βMCD (D = 0.111, p = 0.8807). Data and replicates: n (axons per condition; data from ≥3 independent experiments): Control = 64; EphA3-Fc = 47; βMCD = 58; EphA3-Fc + βMCD = 59; EphA3-Fc + KYL = 52.

Analysis of the cumulative distribution of β deviation angles (Fig. 3D) showed that the EphA3-Fc gradient shifted the distribution rightward, toward positive values. Addition of βMCD to the EphA3-Fc gradient produced an intermediate distribution that did not differ significantly from either the controls or the EphA3-Fc gradient alone. Addition of KYL to the EphA3-Fc gradient produced a distribution that did not differ from the control (Fig. 3F).

The rose diagram of axonal directions (Fig. 3H) showed that EphA3-Fc-exposed nasal RGC axons were concentrated mainly between +20° and +30°, with additional axons distributed between -20° and +50°, consistent with a positive chemotactic pattern. Under control conditions, most axons were oriented between -20° and 0°, whereas βMCD alone produced a peak between 0° and +15°; both distributions were considered neutral. With EphA3-Fc + βMCD, the highest density remained between +20° and +30°, but the broader distribution was consistent with a moderate-to-mild attractive pattern. With EphA3-Fc + KYL, the distribution was neutral, with the highest density between +10° and +20° and lower densities at more extreme angles (Fig. 3H).

Analysis of β deviation angles relative to the initial angles α (Fig. 3I) showed that the EphA3-Fc gradient significantly shifted nasal RGC axons toward positive values (Kolmogorov-Smirnov test, p = 0.0016). In contrast, the two non-gradient controls, with and without βMCD, showed distributions centered around 0 and did not differ significantly from each other. EphA3-Fc + βMCD produced an intermediate distribution that did not differ significantly from either the controls or EphA3-Fc alone. EphA3-Fc + KYL did not differ significantly from the controls but differed significantly from the EphA3-Fc gradient.

Together, these results show that the EphA3-Fc gradient induces a robust attractive turning of nasal RGC axons. Cholesterol depletion with βMCD does not alter basal orientation of nasal RGC axons, indicating that the mere alteration of membrane fluidity is insufficient to redirect axonal growth. However, cholesterol depletion via βMCD diminishes the intensity of the attractive effect exerted by the EphA3-Fc gradient. This suggests that the efficiency of EphA3 signaling depends, at least in part, on cholesterol-enriched domains (lipid rafts), indicating that EphA3-Fc-induced attraction is attenuated in the absence of cholesterol. Moreover, KYL-mediated inhibition of axonal EphA4 signaling abolishes EphA3-Fc-induced attraction. These findings indicate that EphA3-dependent chemoattraction requires basal or appropriately regulated axonal EphA4 forward signaling.

### 3. Plasma membrane cholesterol depletion and EphA4 inhibition differentially affect EphA3-induced axon growth dynamics

Because biased turning and growth-rate modulation can jointly guide axons (Mortimer et al., 2010), we next examined global growth velocity and the temporal organization of extension, pause, and retraction phases during the 2-h recordings.

Analysis of average axon growth velocity (Fig. 4A) showed that the EphA3 gradient significantly increased this parameter in nasal axons. In contrast, cholesterol depletion with βMCD significantly reduced average velocity relative to the control. EphA3-Fc + βMCD produced a growth rate that was lower than that observed with EphA3-Fc alone, higher than that observed with βMCD alone, and not significantly different from the control.

**Figure 4.**
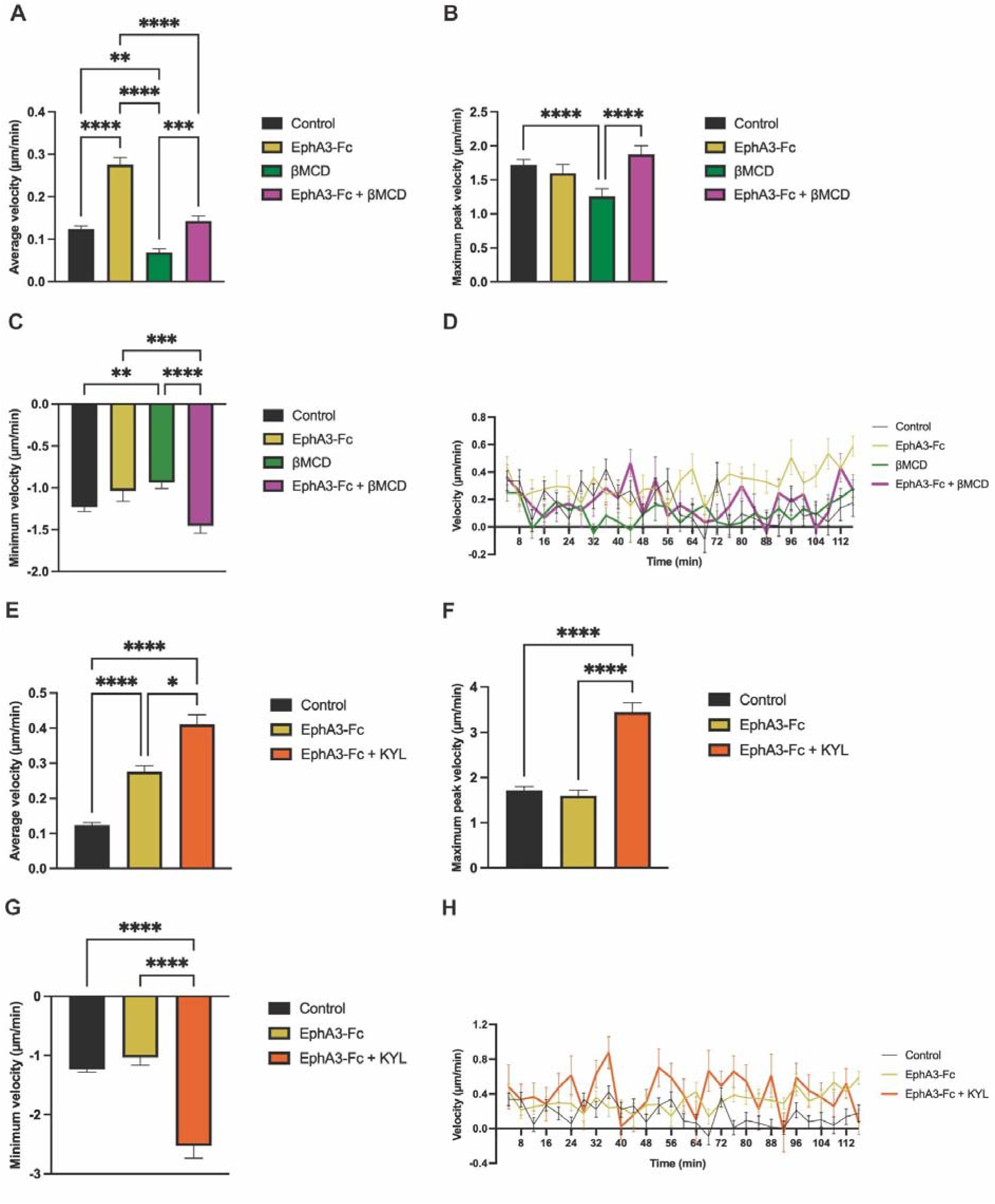
EphA3-Fc–dependent changes in axon growth velocity are modulated by cholesterol depletion and inhibition of EphA activity. Time-lapse recordings were used to quantify axon growth dynamics under the indicated conditions. Velocity was calculated from frame-to-frame growth cone displacement and expressed as µm/min. For each axon, average velocity (mean across the entire recording), maximum peak velocity (average of highest velocity reached per axon), and minimum velocity (average of lowest velocity reached per axon; negative values indicate retraction) were extracted. (A–D) Effects of EphA3-Fc and βMCD, alone or combined. (A) Average velocity. (B) Maximum peak velocity. (C) Minimum velocity. (D) Time-course curves of mean growth speed over 120 min. (E–H) Effects of KYL in the presence of EphA3-Fc gradient. (E) Average velocity. (F) Maximum peak velocity. (G) Minimum velocity. (H) Time-course curves of mean growth speed over 120 min. Bar graphs show mean ± SEM. Differences were assessed using Kruskal–Wallis followed by Dunn’s post hoc multiple-comparisons test. Statistical significance for the indicated pairwise comparisons is shown by brackets and asterisks (* p < 0.05, ** p < 0.01, *** p < 0.001, **** p < 0.0001). n (axons per condition; data from ≥3 independent experiments): Control = 64; EphA3-Fc = 47; βMCD = 58; EphA3-Fc + βMCD = 59; EphA3-Fc + KYL = 52.

Analysis of the maximum peaks in axonal growth rate (Fig. 4B) showed no significant differences among the EphA3-Fc gradient, EphA3-Fc + βMCD, and control conditions, whereas βMCD alone significantly reduced this parameter relative to the other conditions.

Analysis of minimum axonal growth velocity (Fig. 4C) showed that βMCD significantly reduced the magnitude of this parameter relative to the other three conditions, whereas neither the EphA3-Fc gradient nor EphA3-Fc + βMCD differed significantly from the control.

Axonal growth velocity oscillated under all conditions during the 120-min time-lapse recordings, but nasal axons exposed to the EphA3-Fc gradient maintained the highest velocities (Fig. 4D).

Analysis of axonal growth kinetics (Fig. 4D) showed that the EphA3-Fc gradient maintained higher velocities than the control throughout most of the recording. βMCD produced the slowest growth pattern, whereas EphA3-Fc + βMCD produced intermediate kinetics between those observed with EphA3-Fc and βMCD alone, with substantial overlap with the control.

Together, these results demonstrate that the EphA3-Fc gradient increases the average growth velocity of nasal RGC axons and maintains higher velocities throughout the 120-min observation period without increasing maximum velocity. Cholesterol depletion with βMCD reduces growth velocity across the recording period and lowers both maximum and minimum velocity values, producing slower growth with a smaller amplitude of variation. Addition of βMCD to the EphA3-Fc gradient abolishes the effect of EphA3 and restores growth-velocity parameters to control levels.

Analysis of average axonal growth velocity (Fig. 4E) showed that combining the EphA3-Fc gradient with KYL-mediated inhibition of axonal EphA4 activation significantly increased velocity relative to both the control and the EphA3-Fc gradient alone.

Analysis of maximum and minimum axonal growth velocities (Fig. 4F, G) showed that EphA3-Fc + KYL significantly increased both parameters relative to the control and EphA3-Fc conditions, which did not differ significantly from each other.

Analysis of axonal growth kinetics (Fig. 4H) showed that EphA3-Fc + KYL produced higher velocities and greater amplitudes of variation than EphA3-Fc alone or the control.

These results demonstrate that EphA4 inhibition enhances the EphA3-induced increase in axonal growth velocity but alters the growth pattern produced by EphA3 alone. Specifically, EphA3-Fc + KYL increases maximum and minimum velocity values, which are unaffected by EphA3 alone, thereby producing a wider amplitude of velocity variation. These changes are opposite to those produced by βMCD.

#### 3.1. EphA3-dependent stabilization of the axon growth phase is abolished by cholesterol depletion and EphA4 inhibition

Heatmaps of axonal velocity over time showed that velocity values were concentrated mainly within an intermediate range under all conditions (Supplementary Fig. 3). Control axons displayed a relatively uniform pattern throughout the recording, whereas EphA3-Fc increased velocity and the proportion and duration of green-yellow intervals. EphA3-Fc + KYL produced the highest velocities and greatest apparent variability, with alternating positive and negative events. In contrast, βMCD produced the lowest velocities, a predominance of blue intervals, and more events near zero. EphA3-Fc + βMCD resembled the control, with a predominance of intermediate blue-green velocities. These qualitative observations suggest that EphA3-Fc prolongs axonal growth intervals, EphA3-Fc + KYL increases both velocity and its fluctuations, and βMCD reduces velocity and heterogeneity.

Phase maps revealed less frequent phase alternation in EphA3-Fc-exposed nasal axons, with larger blocks of sustained growth (green), shorter retraction phases (red), and a tendency toward shorter pause phases. In contrast, βMCD increased both the frequency and duration of pause phases (yellow). Combining the EphA3-Fc gradient with either βMCD or KYL shortened the prolonged growth episodes and lengthened retraction phases, producing patterns that did not differ significantly from the control. Addition of βMCD tended to abolish the EphA3-induced reduction in pause duration, whereas addition of KYL preserved this reduction (Fig. 5A,B).

**Figure 5.**
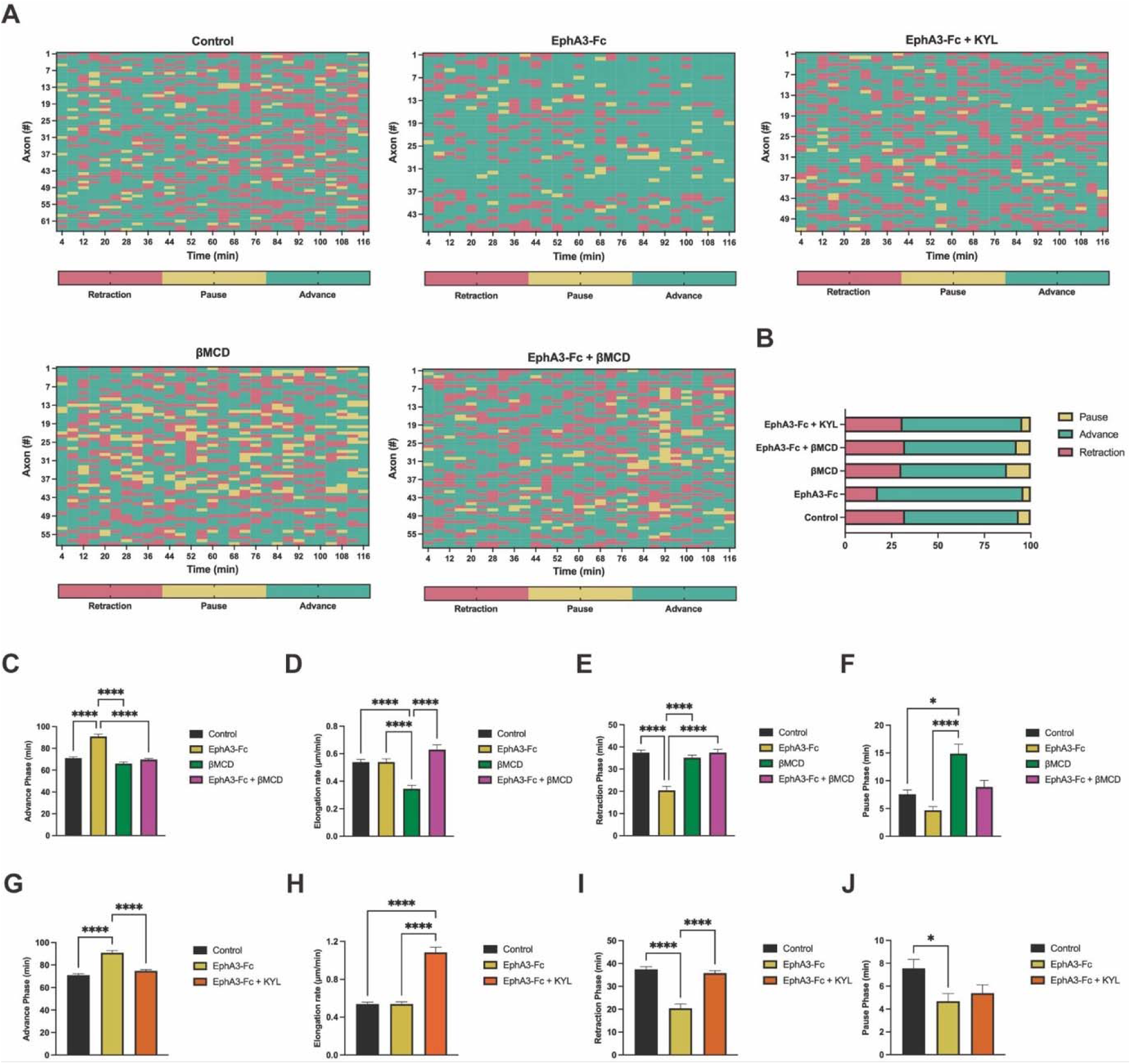
Temporal dynamics of axon growth: The EphA3-dependent increase in stability of axon growth phase is abrogated by cholesterol depletion and inhibition of EphA4 activity. (A) Phase maps of axon growth (per axon and time interval). Each row represents one axon and each column, a 4-min interval. Emerald green = growth, maize yellow = pause, coral = retraction. (B) Overall distribution of axon kinetic states. The stacked bar plot shows the total percentage of time that axons in each condition spend in growth (emerald green), retraction (coral), and pause (maize yellow). Bar graphs showing mean ± SEM of (i) cumulative time spent in growth (C,G), retraction (E,I), and pause (F,J) phases over 2 h and (ii) elongation rate (mean speed during the growth phase) (D,H). Differences were assessed using Kruskal–Wallis followed by Dunn’s post hoc test. Asterisks: *p < 0.05, ****p < 0.0001; ns, not significant. n (axons per condition; data from ≥3 independent experiments): Control = 64; EphA3-Fc = 47; βMCD = 58; EphA3-Fc + βMCD = 59; EphA3-Fc + KYL = 52.

Quantification of the time spent in each phase (Fig. 5C,E,F,G,I,J) and velocity during the growth phase (Fig. 5D,H) showed that the EphA3-Fc gradient increased growth-phase duration and reduced retraction-phase duration without significantly altering elongation rate (Fig. 5C-E). βMCD significantly increased pause-phase duration and reduced elongation rate. EphA3-Fc + βMCD abolished the growth phase changes induced by either treatment alone and restored the βMCD-induced reduction in elongation rate to control levels. Consequently, phase durations and growth velocity during the growth phase (elongation rate) under the combined treatment show no significant differences compared to the control (Fig. 5C-E). Addition of KYL to the EphA3-Fc gradient abolished the increased growth-phase duration and reduced retraction-phase duration induced by EphA3, preserved the trend toward shorter pause duration, and significantly increased elongation rate (Fig. 5G-J).

Overall, EphA3 stabilized axonal advance without increasing elongation rate, whereas βMCD and KYL abolished this stabilization through distinct changes in growth-, pause-, and retraction-phase duration and in elongation rate.

#### 3.2. EphA3 alters the duration of growth, pause, and retraction intervals

Analysis of frequencies of intervals in the different phases of axon growth showed that the extension of intervals of the growth phase (advances) is longer than those of retraction and pause in all conditions (control, EphA3-Fc, βMCD, EphA3-Fc + βMCD and EphA3-Fc + KYL) (Fig. 6).

**Figure 6.**
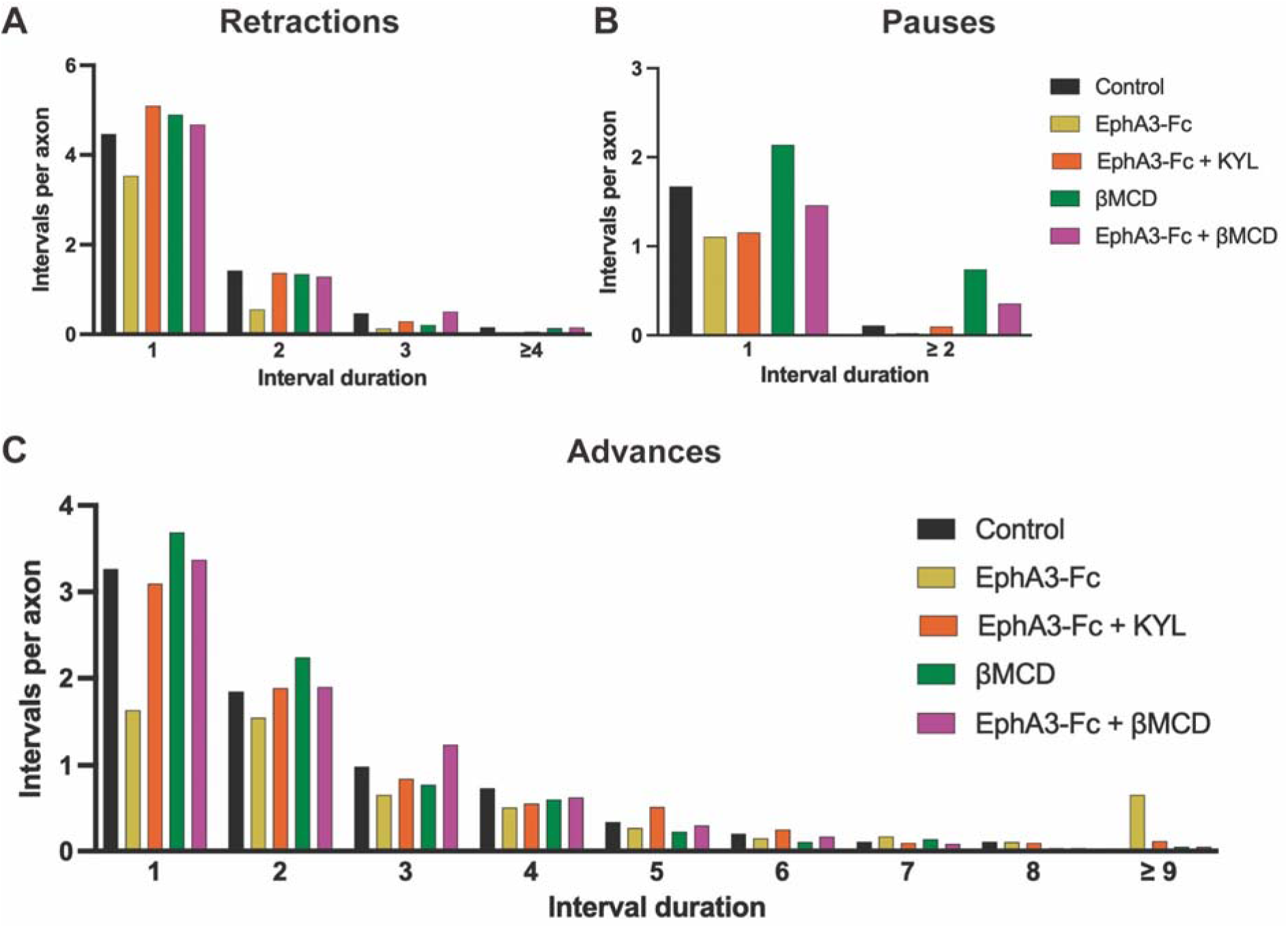
Cholesterol depletion and inhibition of EphA4 activity abrogate the EphA3-dependent increase in the frequency of long intervals of the growth phase and the EphA3-dependent decrease in the frequency of the retractions in nasal RGCs. Distribution of dynamic episode durations in an EphA3-Fc gradient and after cholesterol depletion. All consecutive intervals of retraction (A), pause (B), and growth (C) were counted for each axon over 2 h (4-min bins). Histograms show the mean number of consecutive runs per axon (Y-axis) for each run-length category (X-axis: number of consecutive intervals; e.g., 1 = 4 min, 2 = 8 min, 3 = 12 min, ≥4 = 16 min or longer). n (axons per condition; data from ≥3 independent experiments): Control = 64; EphA3-Fc = 47; βMCD = 58; EphA3-Fc + βMCD = 59; EphA3-Fc + KYL = 52.

Analysis of retraction phase frequencies (Fig. 6A) showed that the EphA3-Fc gradient reduced the frequency of retraction episodes (1–3 intervals) relative to the control. βMCD, EphA3-Fc + βMCD, and EphA3-Fc + KYL reversed this effect, restoring the distribution of retraction-interval durations to control-like levels.

Analysis of pause phase frequencies (Fig. 6B) showed that EphA3-Fc treatment decreases pauses spanning 1 to ≥2 intervals. In contrast, βMCD induces a slight increase in short 1-interval pauses and a more pronounced increase in ≥2-interval pauses. The EphA3-Fc + βMCD co-treatment leads to a slight reduction in short pause intervals (1 interval) and a substantial increase in pauses of ≥2 intervals compared to the control condition, presenting intermediate values between those of EphA3-Fc and βMCD alone. The combination of EphA3-Fc and KYL maintains the reduction in short-interval pauses (1 interval) induced by the EphA3-Fc gradient, but reverses the EphA3-mediated decrease in pauses lasting ≥2 intervals.

Analysis of growth phase frequencies (Fig. 6C) showed that the EphA3-Fc gradient increased the proportion of long episodes (7 and 9 intervals) while reducing short and intermediate ones (1–6 intervals). Cholesterol depletion with βMCD concentrates growth activity within 1–2 interval bouts, reducing intermediate (3–4 intervals) and long (≥5 intervals) episodes. The dual EphA3-Fc + βMCD treatment shows a slight increase in intermediate 3-interval growth episodes, displaying values that are intermediate and similar to the control across short and long intervals. The addition of KYL to the EphA3-Fc gradient reverses the effects of EphA3-Fc on axonal growth phase frequencies.

These patterns confirm that EphA3-Fc prolongs continuous axonal advance, whereas cholesterol depletion fragments growth into shorter episodes. The intermediate pattern produced by EphA3-Fc + βMCD suggests partial compensation between these effects.

Finally, KYL-mediated inhibition of axonal EphA4 activity reversed the EphA3-dependent increase in the duration of growth episodes and the decrease in the duration of retraction episodes, while preserving the smaller EphA3-dependent reduction in pause duration.

#### 3.3. EphA3 reduces transitions among axonal growth phases

EphA3-Fc reduced transitions among growth, pause, and retraction phases, whereas βMCD modestly increased them, indicating opposite effects on temporal stability (Table 1). Adding βMCD or KYL to EphA3-Fc abolished this effect and produced transition frequencies that did not differ from the control.

**Table 1.** Cholesterol depletion and inhibition of EphA4 activity abrogate the EphA3-dependent decrease in the temporal variability of growth phases in nasal RGC axons. The number of phase changes per axon was quantified over recordings of equal duration. Data are shown as mean ± SD. The global comparison across conditions was performed using the Kruskal-Wallis test (Kruskal-Wallis test, H = 48.43, p = 7.68 × 10 ¹ ). P values from pairwise Mann-Whitney U tests were adjusted for multiple comparisons using the Benjamini-Hochberg false discovery rate procedure. The “Interpretation” column indicates whether the number of phase changes per axon was increased or decreased in the second condition relative to the first condition listed in each pairwise comparison. n.s., not significant. Control = 64; EphA3-Fc = 47; βMCD = 58; EphA3-Fc + βMCD = 59; EphA3-Fc + KYL = 52.

| Comparison | Phase changes per axon, mean ± SD | q value | Interpretation |
| --- | --- | --- | --- |
| Control vs EphA3-Fc | 14.9 ± 2.8 vs 10.1 ± 5.0 | $1.17 \times 10^{-6}$ | EphA3-Fc decreased phase switching |
| Control vs βMCD | 14.9 ± 2.8 vs 16.3 ± 3.4 | $3.55 \times 10^{-2}$ | βMCD increased phase switching |
| Control vs EphA3-Fc + βMCD | 14.9 ± 2.8 vs 15.2 ± 3.4 | $5.36 \times 10^{-1}$ | n.s. |
| Control vs EphA3-Fc + KYL | 14.9 ± 2.8 vs 14.5 ± 2.8 | $5.29 \times 10^{-1}$ | n.s. |
| EphA3-Fc vs EphA3-Fc + KYL | 10.1 ± 5.0 vs 14.5 ± 2.8 | $1.48 \times 10^{-5}$ | EphA3-Fc + KYL increased phase switching |
| EphA3-Fc vs EphA3-Fc + βMCD | 10.1 ± 5.0 vs 15.2 ± 3.4 | $1.17 \times 10^{-6}$ | EphA3-Fc + βMCD increased phase switching |
| EphA3-Fc vs βMCD | 10.1 ± 5.0 vs 16.3 ± 3.4 | $2.44 \times 10^{-6}$ | βMCD increased phase switching |
| EphA3-Fc + KYL vs βMCD | 16.3 ± 3.4 vs 14.5 ± 2.8 | $1.03 \times 10^{-2}$ | βMCD increased phase switching |
| βMCD vs EphA3-Fc + βMCD | 16.3 ± 3.4 vs 15.2 ± 3.4 | $1.59 \times 10^{-1}$ | n.s. |
| EphA3-Fc + βMCD vs EphA3-Fc + KYL | 15.2 ± 3.4 vs 14.5 ± 2.8 | $2.52 \times 10^{-1}$ | n.s. |

Growth persistence (1 to 1) was the most frequent transition under every condition, followed by retraction-to-growth (-1 to 1) and growth-to-retraction (1 to -1) transitions (Tables 2 and 3; Fig. 7). EphA3-Fc increased overall stability (persistence in the same phase: 1 to 1 + -1 to -1 + 0 to 0) by 15.9-22.3% relative to all other conditions (all q < 0.001). βMCD reduced stability relative to the control (-5.02, q = 0.0048), whereas EphA3-Fc + βMCD and EphA3-Fc + KYL did not differ much from the control.

**Figure 7.**
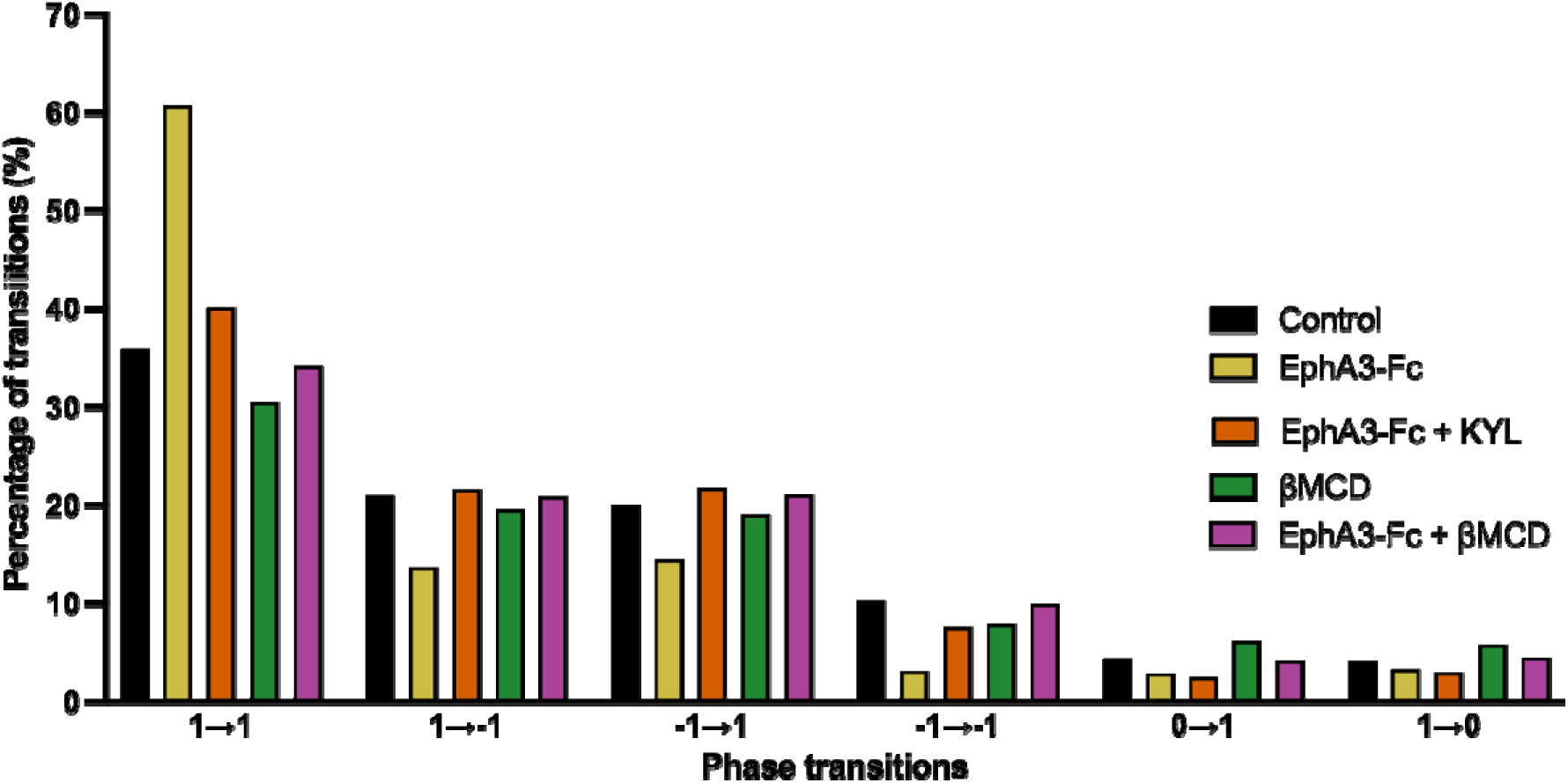
Cholesterol depletion and inhibition of EphA4 activity abrogate the EphA3-dependent decrease in the temporal variability of growth phases in nasal RGC axons. Bar graph represents the transitions between axonal growth phases under control, EphA3-Fc, βMCD, EphA3-Fc + βMCD and EphA3-Fc + KYL conditions. Comparison of the percentage (%) of phase transitions per condition. Each bar group corresponds to a transition type: 1→1 (growth persistence), 1→−1 (growth→retraction), −1→1 (retraction→growth), −1→−1 (retraction persistence), 0→1 (pause→growth), and 1→0 (growth→pause). Bar height represents the percentage of events relative to the total number of transitions within each condition (Control, EphA3-Fc, βMCD, EphA3-Fc + βMCD, and EphA3-Fc + KYL). Control = 64; EphA3-Fc = 47; βMCD = 58; EphA3-Fc + βMCD = 59; EphA3-Fc + KYL = 52.

**Table 2.** Cholesterol depletion and inhibition of EphA4 activity abrogate the EphA3-dependent increase in the stability of the growth phase in nasal RGC axons. The percentages of the 6 types of transitions are represented in each one of the analyzed conditions (control, EphA3-Fc, βMCD, EphA3-Fc + βMCD and EphA3-Fc + KYL). Persistence in growth: 1→1, persistence in retraction: -1→-1, Growth to retraction: 1→-1, Retraction to growth: -1→1, Pause to growth: 0→1, Growth to pause: 1→0. The stability column includes the percentage of time without any change of phase (1→1, -1→-1, 0→0). Control = 64; EphA3-Fc = 47; βMCD = 58; EphA3-Fc + βMCD = 59; EphA3-Fc + KYL = 52.

| Condition | 1→1 | 1→-1 | -1→1 | -1→-1 | 0→1 | 1→0 | Stability |
| --- | --- | --- | --- | --- | --- | --- | --- |
| Control | 36% | 21% | 20.1% | 10.3% | 4.4% | 4.1% | 46.7% |
| EphA3-Fc | 60.7 % | 13.7% | 14.5% | 3.1% | 2.9% | 3.3% | 64% |
| EphA3-Fc + KYL | 40.2% | 21.6% | 21.8% | 7.6% | 2.5% | 3% | 48.1% |
| βMCD | 30.5% | 19.6% | 19.1% | 7.9% | 6.2% | 5.8% | 41.7% |
| EphA3-Fc + βMCD | 34.2% | 20.9% | 21.1% | 10% | 4.2% | 4.5% | 45.6% |

**Table 3.**
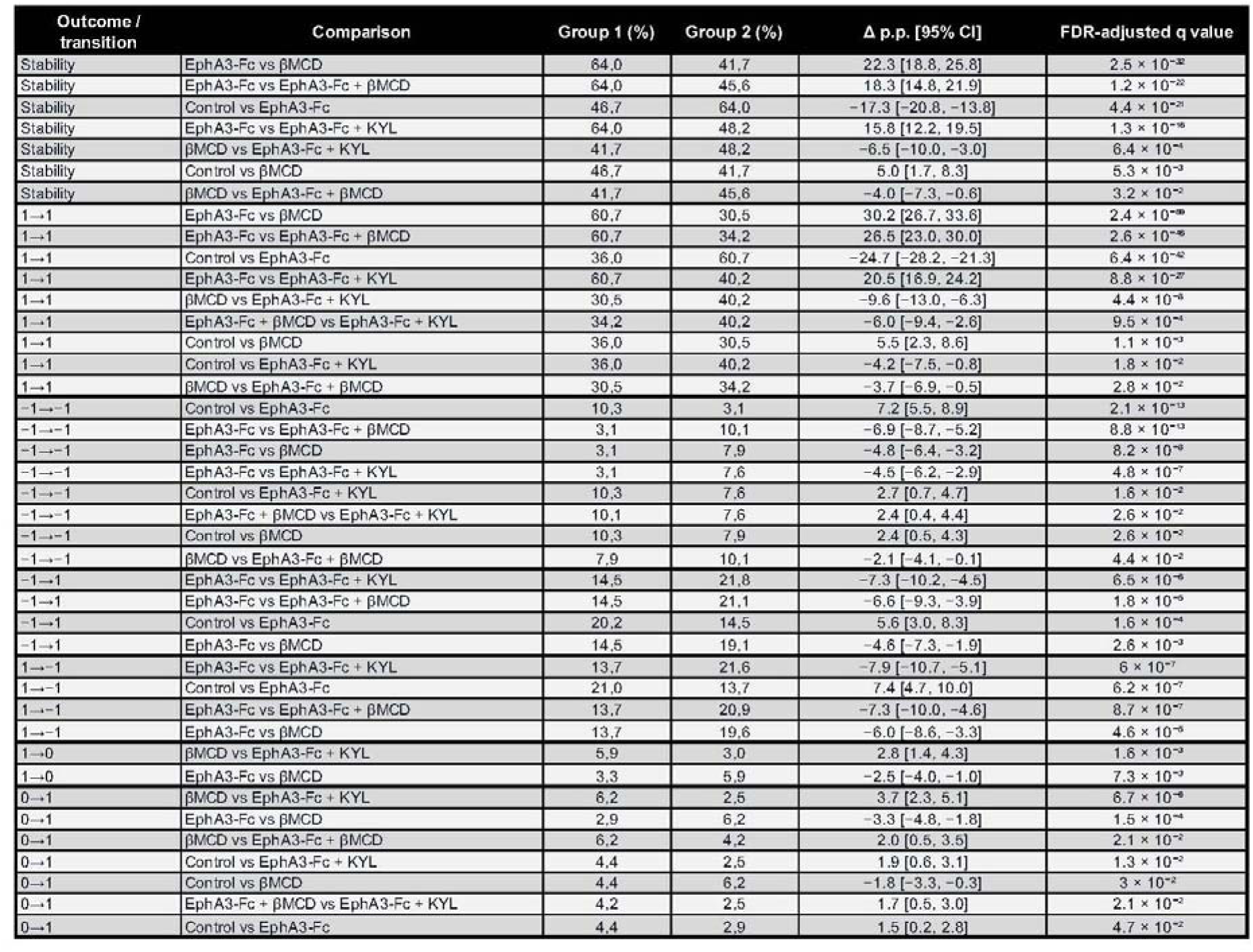
Cholesterol depletion and inhibition of EphA4 activity abrogate the EphA3-dependent increase in the stability of the growth phase and the EphA3-dependent decrease in the temporal variability of growth phases in nasal RGC axons. Pairwise comparisons of the proportions of the different transition types for the Control, EphA3-Fc, βMCD, EphA3-Fc + βMCD and EphA3-Fc + KYL conditions are represented. Only pairwise comparisons that remained statistically significant after false discovery rate correction are shown. For each outcome or transition category, percentages indicate the proportion of events observed in each experimental group. In the Comparison column, the condition listed before “vs” corresponds to Group 1, whereas the condition listed after “vs” corresponds to Group 2. Differences are expressed as percentage points and were calculated as the percentage in Group 1 minus the percentage in Group 2; therefore, negative values indicate a higher percentage in Group 2, whereas positive values indicate a higher percentage in Group 1. The 95% confidence interval refers to the difference in percentage points between groups. FDR-adjusted q values are reported, with q < 0.05 considered statistically significant. Transition categories are indicated as initial state → final state; the definitions of states 1, −1, and 0 are described in the Methods. NR, nasal retina; TR, temporal retina; CI, confidence interval; FDR, false discovery rate. The complete set of pairwise comparisons, including non-significant results and raw counts, is provided in Supplementary Table 1.

The strongest effect involved growth persistence: EphA3-Fc increased 1-to-1 transitions by 20.5-30.2% relative to the other conditions (all q < 0.001). βMCD moderately reduced growth persistence relative to the control, whereas both combined treatments restored it to control-like values that remained higher than those observed with βMCD alone (Tables 2 and 3; Fig. 7).

EphA3-Fc reduced retraction persistence relative to every other condition and markedly reduced growth-to-retraction transitions. βMCD and EphA3-Fc + KYL also modestly reduced retraction persistence relative to the control, whereas EphA3-Fc + βMCD did not. Retraction-to-growth transitions were less frequent with EphA3-Fc than under all other conditions, consistent with fewer retraction episodes requiring recovery.

βMCD increased pause-to-growth transitions relative to all other conditions and increased growth-to-pause transitions relative to EphA3-Fc and EphA3-Fc + KYL. EphA3-Fc + βMCD produced intermediate, control-like values. Detailed pairwise comparisons are reported in Tables 2 and 3 and Supplementary Table 1.

Collectively, EphA3-Fc-induced attraction and sustained growth required plasma membrane cholesterol: βMCD attenuated turning, reduced basal elongation rate, increased pausing and phase switching, and abolished EphA3-Fc-induced growth stability. Basal EphA4 activity was also required for turning. KYL abolished attraction and phase stabilization but increased average and maximum velocity, dissociating directional guidance and growth persistence from intrinsic elongation rate.

### 4. Integrated analysis of axonal guidance and growth

Turning angle β showed no consistent association with initial orientation, initial axon length, position within the gradient, average velocity, growth persistence, or pause fraction. Initial orientation and gradient position were also unrelated to average velocity. The only condition-specific association was a negative correlation between initial axon length and average velocity under the EphA3-Fc + KYL condition (Spearman’s ρ = -0.53, p < 0.0001). Thus, the measured baseline characteristics and growth dynamics did not consistently explain guidance responses, although smaller or condition-specific effects remain possible (Supplementary Results; Supplementary Figs. 4-12).

## DISCUSSION

This study shows that EphA/ephrin-A guidance comprises separable regulation of directional turning and net axon extension. Clustered EphA3 acted as both a haptotactic and chemoattractive cue for nasal RGC axons. In a gradient, it promoted turning toward higher concentrations and increased average axon growth velocity by prolonging stable growth and reducing retraction, without changing maximum elongation velocity. Guidance cues therefore generate kinetic profiles that cannot be inferred from final growth cone position alone.

### Directional turning and growth dynamics are independently regulated

Molecular gradients more closely approximate the environment encountered *in vivo* than stripe assays at fixed concentrations. Combined with time-lapse imaging, they allowed us to distinguish direction from net extension and to resolve extension into phase persistence and elongation rate. Models predict that biased turning and growth-rate modulation can contribute jointly to guidance, depending on gradient steepness (Mortimer et al., 2010). This interpretation is consistent with models in which extracellular signals regulate average axon growth by altering transitions between growing and paused states (Padmanabhan and Goodhill, 2018). Our findings demonstrate both mechanisms take place within one experimental system.

Axonal turning did not correlate consistently with average axon growth velocity, growth persistence, pausing, initial axon length, or gradient orientation. This dissociation is compatible with localized second-messenger and cytoskeletal signaling controlling direction while more global pathways regulate elongation (Akiyama and Kamiguchi, 2010; Myers et al., 2011). Local protein synthesis contributes to growth-cone turning responses to netrin-1 and Sema3A (Campbell and Holt, 2001; Leung et al., 2006). Guidance cues can also regulate distinct aspects of neuronal morphology separately: netrin-1 and Sema3A alter cortical axon branching without significantly changing axon length (Dent et al., 2004). EphA3 increased net extension by stabilizing advance rather than raising maximum velocity, consistent with pauses and retractions constraining RGC growth capacity (Steketee et al., 2014). This suggests that phase sequence and duration constitute a dynamic code for interpreting positional information. These findings explain why similar net growth can arise from distinct combinations of persistence and instantaneous elongation.

### EphA3 generates a distinct guidance program

Although EphA3 is generally considered a repulsive receptor, its ectodomain acted as a ligand that attracted nasal RGC axons and promoted extension. This extends previous evidence that tectal EphA3 stimulates nasal RGC growth toward the caudal tectum and prevents premature branching and termination rostrally (Ortalli et al., 2012; Fiore et al., 2019; Medori et al., 2020). Our dynamic analysis indicates that EphA3 promotes positive turning and stabilizes growth while reducing retractions.

### Plasma membrane cholesterol supports basal growth and EphA3 signaling

Cholesterol depletion reduces basal axonal average velocity by lowering elongation rate and growth-phase stability while increasing pauses and phase transitions. It also attenuated EphA3-induced attraction and abolished sustained growth stimulation. This dependence is consistent with ephrin-As and their co-receptors occupying cholesterol-rich lipid rafts. These membrane microdomains may enable cis interactions and couple Eph/ephrin complexes to Src-family kinases, integrins, Rho GTPases, and cytoskeletal regulators (Bonanomi et al., 2012; Hernaiz-Llorens et al., 2021). Their disruption could destabilize conversion of EphA3 binding into polarized, persistent growth (Cecchini et al., 2022). ADAM10-mediated ephrin-A cleavage provides another membrane-dependent regulatory step (Sullivan et al., 2023). The variable effects of βMCD across neuronal systems likely reflect differences in cell type, basal cholesterol concentration, treatment, and signaling context (Guirland et al., 2004; Tassew et al., 2014; Roselló-Busquets et al., 2019; Griswold et al., 2025).

### EphA4 modulation is required for EphA3-dependent attraction

KYL inhibition of ephrin-A-dependent EphA4 forward signaling abolished EphA3-induced axon attraction but increased and altered its growth-promoting effect: net growth increased through a higher elongation rate rather than greater phase stability. Basal EphA4 activity is therefore required for the directional effect of EphA3, further demonstrating the independent regulation of turning, growth persistence, and elongation rate.

We propose that basal axonal EphA4 activity enables tectal EphA3 to promote nasal axon growth and guidance rostrally. EphA3 may compete with axonal EphA4 for ephrin-A binding, reducing canonical forward signaling and favoring extension. More caudally in the tectum, declining EphA3 and increasing ephrin-A would raise EphA4 activity, slow growth, and permit branching and termination. This model is compatible with EphA/ephrin-A coadaptation through dynamic cis interactions and membrane redistribution (Fiederling et al., 2017; Medori et al., 2020). Furthermore, sustained EphA4 inhibition may promote regeneration (Chen et al., 2022; Ning et al., 2020; Verma et al., 2023) but could impair accurate target mapping (Fig. 8).

**Figure 8.**
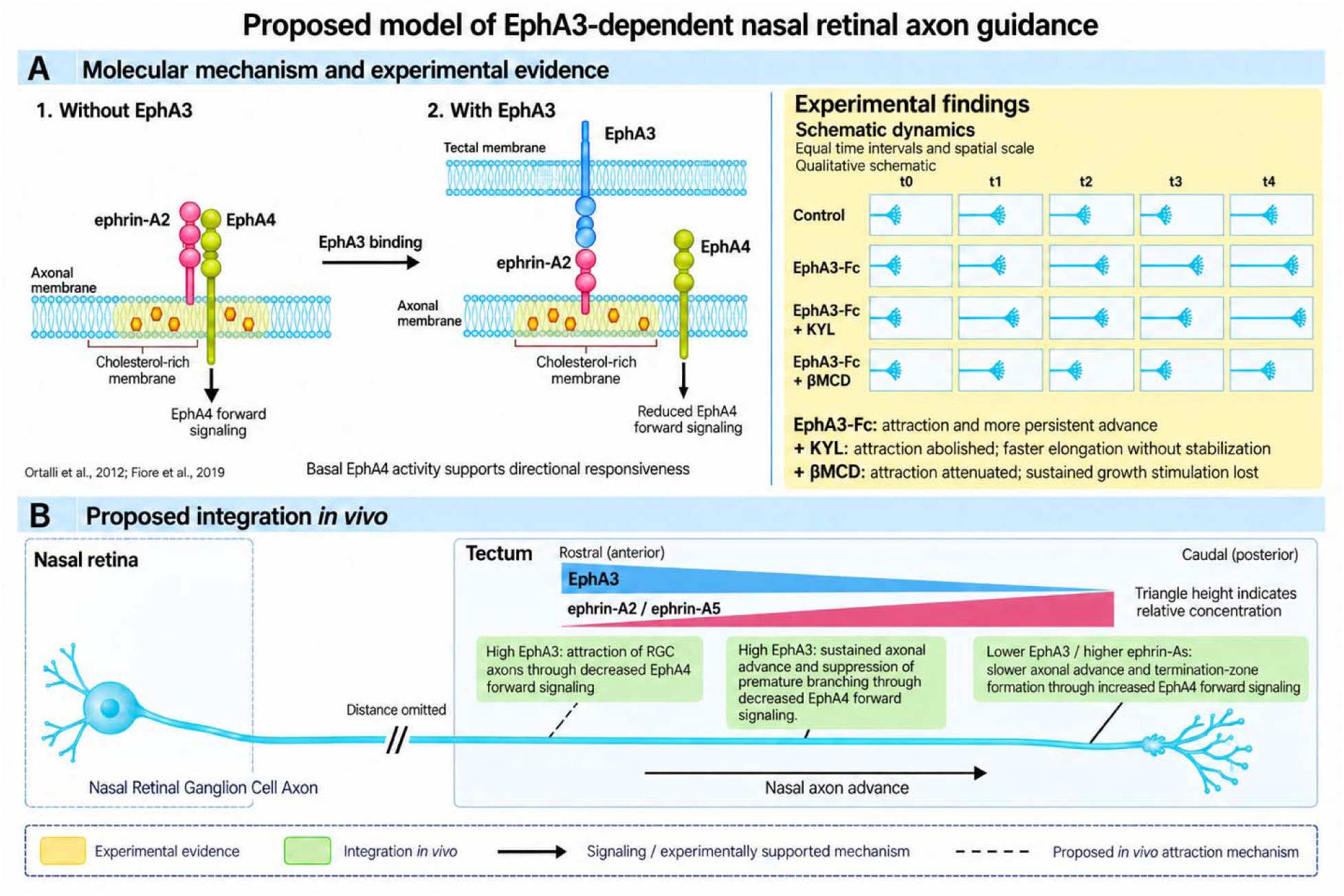
Proposed model of EphA3-dependent nasal retinal axon guidance. (A) Two molecular states illustrate the modulation of axonal EphA4 forward signaling. In the absence of EphA3, axonal ephrin-A2 associates with EphA4 and supports its forward signaling. Tectal EphA3 interacts with axonal ephrin-As and decreases EphA4 forward signaling through competition and membrane redistribution (Ortalli et al., 2012; Fiore et al., 2019). Cholesterol-dependent membrane organization may support these interactions. *In vitro*, clustered EphA3-Fc attracts nasal retinal ganglion cell (RGC) axons and stabilizes advance by prolonging growth episodes and reducing retractions and phase transitions. In the presence of EphA3-Fc, EphA4 inhibition with KYL abolishes attraction and disrupts growth stabilization while increasing the elongation rate. Cholesterol depletion with βMCD attenuates attraction and abolishes sustained growth stimulation. These perturbations support a requirement for EphA4 signaling and membrane cholesterol for tectal EphA3 actions on nasal RGC axonal growth and guidance. (B) Proposed integration along the rostrocaudal tectal axis. High rostral EphA3 attracts nasal RGC axons through reduced EphA4 forward signaling, favoring tectal invasion. EphA3 also promotes sustained axonal advance and suppresses premature branching through decreased EphA4 forward signaling. As axons extend caudally, declining EphA3 and increasing ephrin-A2/ephrin-A5 are proposed to increase EphA4 forward signaling, slow axonal advance, and permit branching and termination-zone formation (Monschau et al., 1997; Yates et al., 2001; Sakurai et al., 2002; Hansen et al., 2004; Ortalli et al., 2012; Fiore et al., 2019). Yellow identifies experimental evidence from the present and previous studies, whereas green identifies the integration of these findings into the proposed *in vivo* model. Solid arrows in A depict signaling and experimentally supported mechanisms. In B, the dashed connection denotes the proposed mechanism of attraction *in vivo*, whereas solid leader lines link the other annotations to the corresponding axonal regions. Molecular shapes, positions, and concentration profiles are schematic.

Accordingly, EphA4 signaling should not be viewed simply as a brake on axon growth. Its basal activity may provide directional competence during pathfinding, whereas stronger forward signaling may restrict extension and favor target-zone formation. This distinction may be particularly relevant when considering EphA4 inhibition as a regenerative strategy.

### Implications for retinotectal mapping

Together, EphA3, ephrin-As, axonal EphA4 activity, and membrane cholesterol can coordinate retinal axon advance and positioning. High rostral tectal EphA3 would attract nasal axons, stabilize caudal growth, and suppress premature branching. As nasal axons advance, increasing ephrin-A2/ephrin-A5 and decreasing EphA3 would slow growth and enable termination-zone formation (Scicolone et al., 2009; Ortalli et al., 2012; Fiore et al., 2019; Medori et al., 2020) (Fig. 8).

In conclusion, EphA/ephrin-A signals regulate directional bias, growth-phase persistence, and elongation rate as distinct but coordinated components. EphA3 attracts nasal RGC axons and stabilizes advance, while its directional and kinetic effects require basal EphA4 activity and cholesterol-dependent membrane organization. This framework helps explain how growth cones integrate positional signals and highlights the need for regenerative strategies to promote extension without sacrificing guidance precision.

