## Supplementary Data for "EphA3 couples attractive guidance to persistent retinal axon growth through modulation of EphA4 forward signaling in a plasma membrane cholesterol-dependent manner"

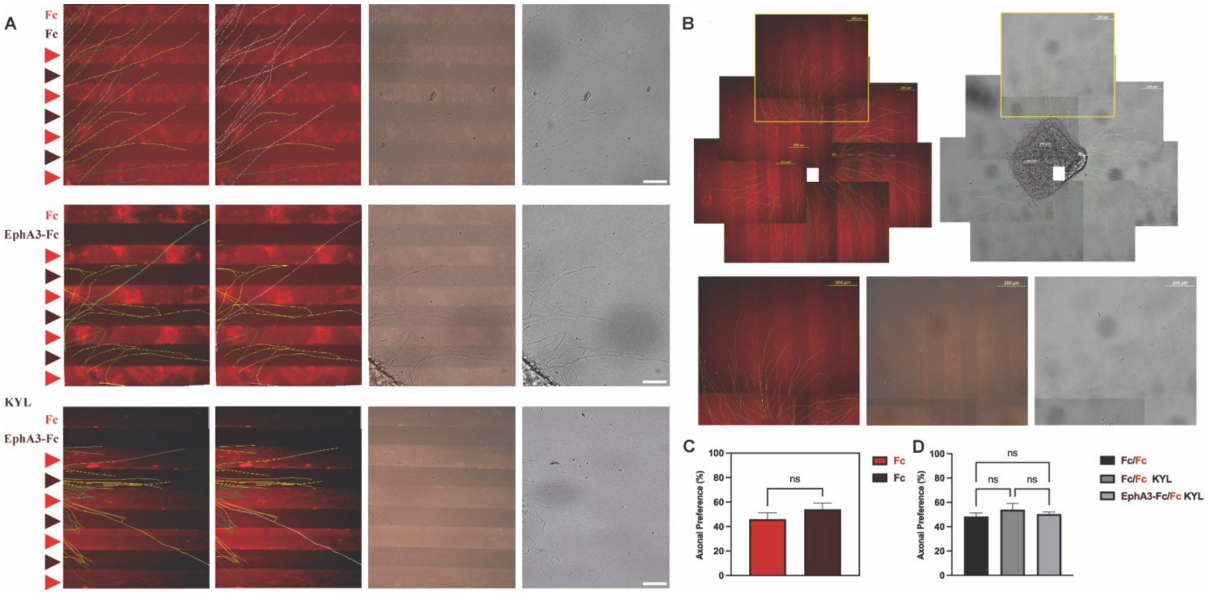


**Supplementary Figure 1. Analysis of axonal growth and EphA-dependent guidance in a stripe substrate choice assay.** (A) Representative images of nasal RGC axon outgrowth on alternating stripe substrates containing unlabeled anti-Fc and Alexa 594–labeled anti-Fc (red; control, top), or clustered EphA3-Fc and Alexa 594–labeled anti-Fc in the absence (middle) or presence (bottom) of KYL. Labeled and unlabeled stripes are indicated by red and dark triangles, respectively. Explants are located to the left of each field. From left to right, images show color-classified axonal trajectories, classified trajectories including axons crossing two or more stripes in white, merged stripe fluorescence and DIC images, and DIC images of the same fields. Green indicates axons terminating on Alexa 594–labeled anti-Fc stripes; yellow indicates axons terminating on clustered EphA3-Fc or unlabeled anti-Fc stripes, as applicable; and white indicates axons crossing two or more stripes. Scale bars: 50 µm. (B) Representative images of a nasal retinal explant growing on alternating stripes of unlabeled anti-Fc and Alexa 594–labeled anti-Fc (red) in the presence of the EphA4-blocking peptide KYL. Reconstructions of epifluorescence (left) and differential interference contrast (DIC; right) images show the explant and axonal trajectories. Axons are classified according to their terminal location: green, axons terminating on Alexa 594–labeled anti-Fc stripes; yellow, axons terminating on unlabeled anti-Fc stripes; and white, axons crossing two or more stripes, indicating no preference. Yellow boxes indicate the distal axonal region shown at higher magnification below as epifluorescence, merged epifluorescence/DIC, and DIC images. Scale bars: 200 µm. (C) Percentage of axons whose growth cones were located on Alexa 594–labeled versus unlabeled anti-Fc stripes in the presence of KYL (Mann–Whitney U test). (D) Percentage of axons located on the unlabeled anti-Fc stripes in control assays, on the unlabeled anti-Fc stripes in control assays performed with KYL, or on clustered EphA3-Fc stripes in EphA3-Fc/KYL assays (Kruskal–Wallis test followed by Dunn's post hoc test). Data are presented as mean ± SEM; ns, not significant.

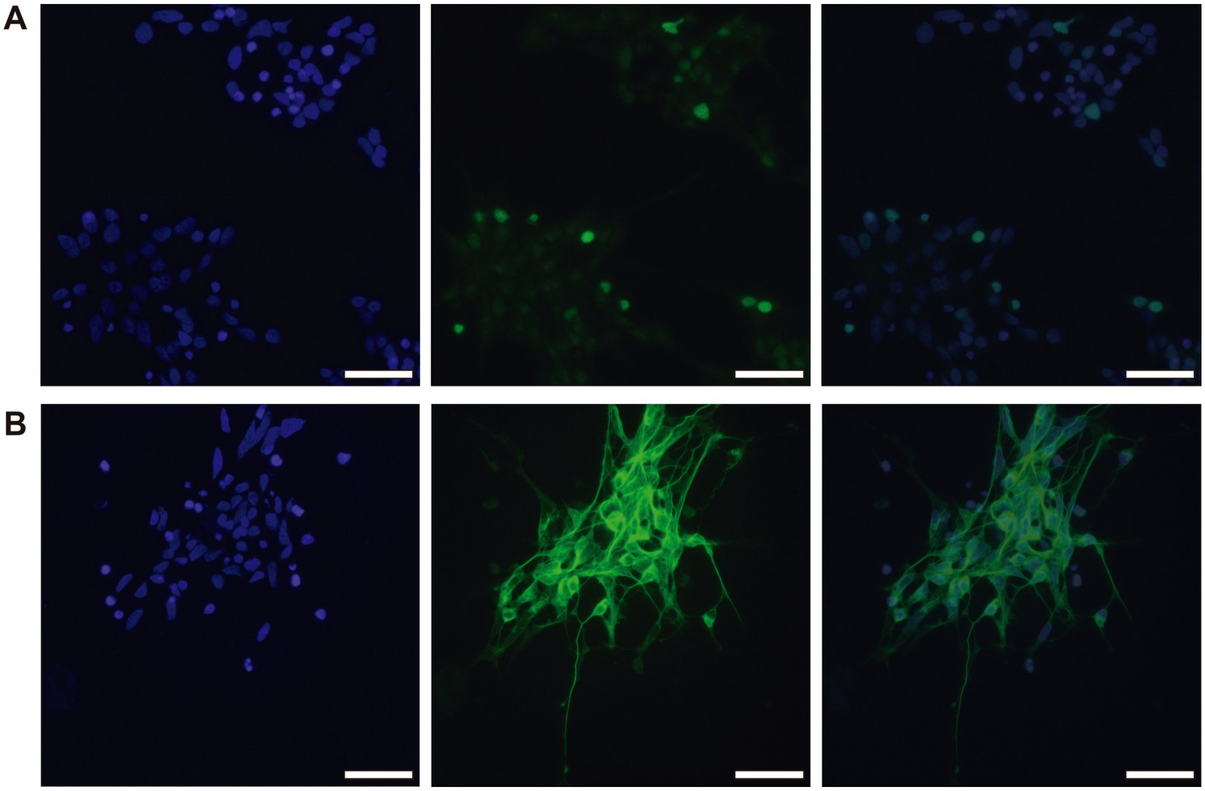


**Supplementary Figure 2. Identification of RGCs in dissociated cultures obtained from chicken embryo retinas.** (A) Immunofluorescence against Islet-2 (green) and nuclear label with Hoechst (blue). (B) Immunofluorescence against βIII-tubulin (neuron-specific, green) and nuclear label with Hoechst (blue). The left column shows the blue channel (Hoechst), the central column shows the green channel (Islet-2 in A and βIII-tubulin in B) and the right column shows the merged image. In order to evaluate only the RGCs, which constitute the only type of retinal projection neuron in this phase of development, only neurons with axons longer than twice the diameter of soma were considered (Ortalli et al., 2012). The identity of RGCs was confirmed by the expression of Islet-2. Cultures were obtained from E7 (HH30-31) chicken embryo retinas. Scale bars = 100 μm.

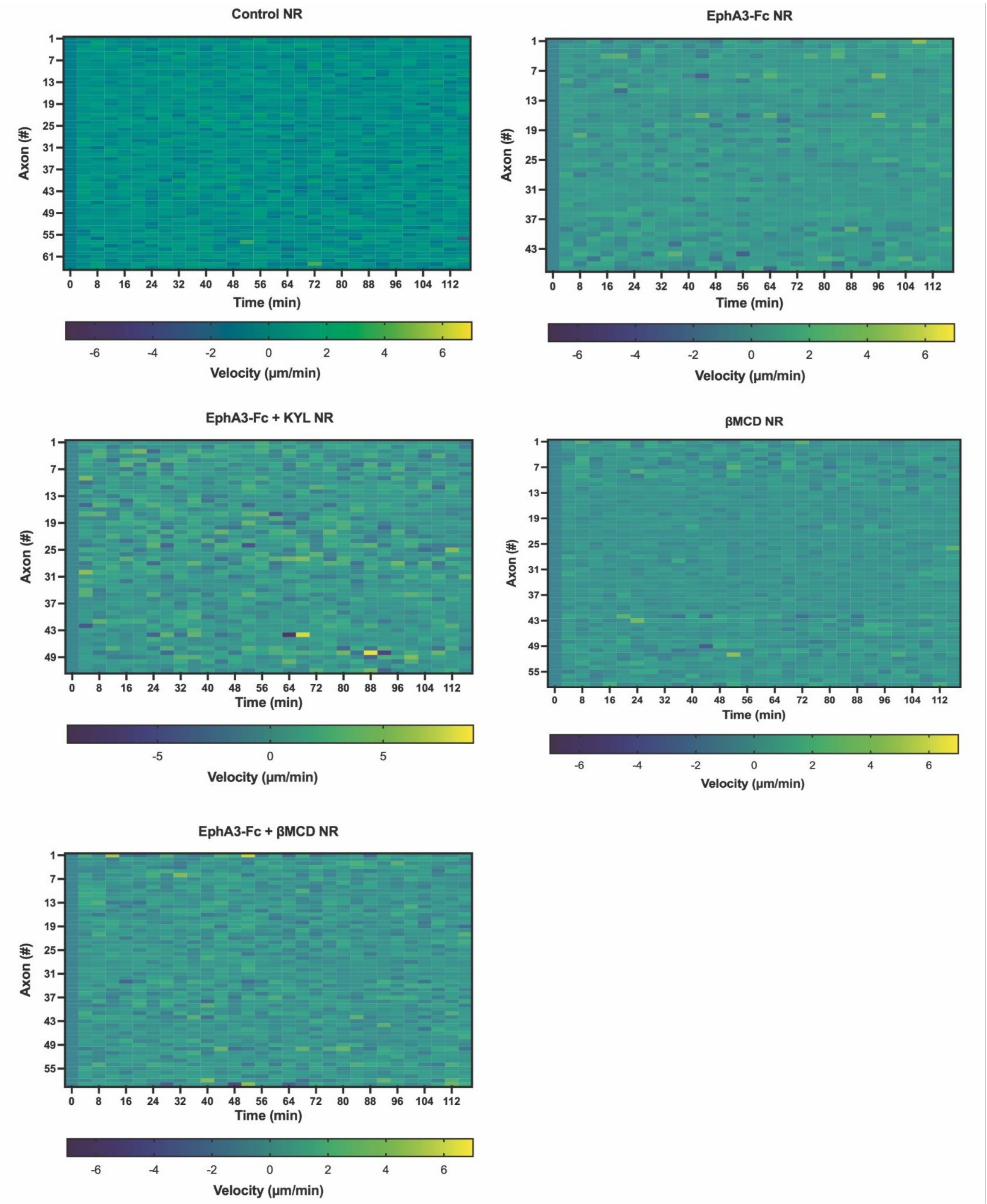


**Supplementary Figure 3. Heat maps of individual axon growth velocities along the time-lapse** of nasal RGC axons exposed to EphA3-Fc (EphA3-Fc NR), EphA3-Fc + KYL (EphA3-Fc + KYL NR), EphA3-Fc + βMCD (EphA3-Fc + βMCD NR) gradients, with βMCD (βMCD NR), and without molecular gradient (Control NR). Each row corresponds to an individual axon and each column represents a 4 min interval along the 120 min of time-lapse. Colours indicate velocity (μm/min).

| Outcome / transition | Comparison | Group 1 (%) | Group 2 (%) | Δ p.p. [95% CI] | FDR-adjusted q value |
|---|---|---|---|---|---|
| Stability | EphA3-Fc vs βMCD | 64,0 | 41,7 | 22.3 [18.8, 25.8] | $2.5 \times 10^{-32}$ |
| Stability | EphA3-Fc vs EphA3-Fc + βMCD | 64,0 | 45,6 | 18.3 [14.8, 21.9] | $1.2 \times 10^{-22}$ |
| Stability | Control vs EphA3-Fc | 46,7 | 64,0 | −17.3 [−20.8, −13.8] | $4.4 \times 10^{-21}$ |
| Stability | EphA3-Fc vs EphA3-Fc + KYL | 64,0 | 48,2 | 15.8 [12.2, 19.5] | $1.3 \times 10^{-16}$ |
| Stability | βMCD vs EphA3-Fc + KYL | 41,7 | 48,2 | −6.5 [−10.0, −3.0] | $6.4 \times 10^{-4}$ |
| Stability | Control vs βMCD | 46,7 | 41,7 | 5.0 [1.7, 8.3] | $5.3 \times 10^{-3}$ |
| Stability | βMCD vs EphA3-Fc + βMCD | 41,7 | 45,6 | −4.0 [−7.3, −0.6] | $3.2 \times 10^{-2}$ |
| Stability | EphA3-Fc + βMCD vs EphA3-Fc + KYL | 45,6 | 48,2 | −2.5 [−6.0, 1.0] | $2 \times 10^{-1}$ |
| Stability | Control vs EphA3-Fc + KYL | 46,7 | 48,2 | −1.4 [−4.9, 2.0] | $4.6 \times 10^{-1}$ |
| Stability | Control vs EphA3-Fc + βMCD | 46,7 | 45,6 | 1.1 [−2.3, 4.4] | $5.3 \times 10^{-1}$ |
| 1→1 | EphA3-Fc vs βMCD | 60,7 | 30,5 | 30.2 [26.7, 33.6] | $2.4 \times 10^{-59}$ |
| 1→1 | EphA3-Fc vs EphA3-Fc + βMCD | 60,7 | 34,2 | 26.5 [23.0, 30.0] | $2.6 \times 10^{-46}$ |
| 1→1 | Control vs EphA3-Fc | 36,0 | 60,7 | −24.7 [−28.2, −21.3] | $6.4 \times 10^{-42}$ |
| 1→1 | EphA3-Fc vs EphA3-Fc + KYL | 60,7 | 40,2 | 20.5 [16.9, 24.2] | $8.8 \times 10^{-27}$ |
| 1→1 | βMCD vs EphA3-Fc + KYL | 30,5 | 40,2 | −9.6 [−13.0, −6.3] | $4.4 \times 10^{-8}$ |
| 1→1 | EphA3-Fc + βMCD vs EphA3-Fc + KYL | 34,2 | 40,2 | −6.0 [−9.4, −2.6] | $9.5 \times 10^{-4}$ |
| 1→1 | Control vs βMCD | 36,0 | 30,5 | 5.5 [2.3, 8.6] | $1.1 \times 10^{-3}$ |
| 1→1 | Control vs EphA3-Fc + KYL | 36,0 | 40,2 | −4.2 [−7.5, −0.8] | $1.8 \times 10^{-2}$ |
| 1→1 | βMCD vs EphA3-Fc + βMCD | 30,5 | 34,2 | −3.7 [−6.9, −0.5] | $2.8 \times 10^{-2}$ |
| 1→1 | Control vs EphA3-Fc + βMCD | 36,0 | 34,2 | 1.8 [−1.4, 5.0] | $2.7 \times 10^{-1}$ |
| −1→−1 | Control vs EphA3-Fc | 10,3 | 3,1 | 7.2 [5.5, 8.9] | $2.1 \times 10^{-13}$ |
| −1→−1 | EphA3-Fc vs EphA3-Fc + βMCD | 3,1 | 10,1 | −6.9 [−8.7, −5.2] | $8.8 \times 10^{-13}$ |
| −1→−1 | EphA3-Fc vs βMCD | 3,1 | 7,9 | −4.8 [−6.4, −3.2] | $8.2 \times 10^{-8}$ |
| −1→−1 | EphA3-Fc vs EphA3-Fc + KYL | 3,1 | 7,6 | −4.5 [−6.2, −2.9] | $4.8 \times 10^{-7}$ |
| −1→−1 | Control vs EphA3-Fc + KYL | 10,3 | 7,6 | 2.7 [0.7, 4.7] | $1.6 \times 10^{-2}$ |
| −1→−1 | EphA3-Fc + βMCD vs EphA3-Fc + KYL | 10,1 | 7,6 | 2.4 [0.4, 4.4] | $2.6 \times 10^{-2}$ |
| −1→−1 | Control vs βMCD | 10,3 | 7,9 | 2.4 [0.5, 4.3] | $2.6 \times 10^{-2}$ |
| −1→−1 | βMCD vs EphA3-Fc + βMCD | 7,9 | 10,1 | −2.1 [−4.1, −0.1] | $4.4 \times 10^{-2}$ |
| −1→−1 | βMCD vs EphA3-Fc + KYL | 7,9 | 7,6 | 0.3 [−1.6, 2.2] | $7.9 \times 10^{-1}$ |
| −1→−1 | Control vs EphA3-Fc + βMCD | 10,3 | 10,1 | 0.3 [−1.8, 2.3] | $7.9 \times 10^{-1}$ |
| −1→1 | EphA3-Fc vs EphA3-Fc + KYL | 14,5 | 21,8 | −7.3 [−10.2, −4.5] | $6.5 \times 10^{-6}$ |
| −1→1 | EphA3-Fc vs EphA3-Fc + βMCD | 14,5 | 21,1 | −6.6 [−9.3, −3.9] | $1.8 \times 10^{-5}$ |
| −1→1 | Control vs EphA3-Fc | 20,2 | 14,5 | 5.6 [3.0, 8.3] | $1.6 \times 10^{-4}$ |
| −1→1 | EphA3-Fc vs βMCD | 14,5 | 19,1 | −4.6 [−7.3, −1.9] | $2.6 \times 10^{-3}$ |
| −1→1 | βMCD vs EphA3-Fc + KYL | 19,1 | 21,8 | −2.8 [−5.6, 0.1] | $1.2 \times 10^{-1}$ |
| −1→1 | βMCD vs EphA3-Fc + βMCD | 19,1 | 21,1 | −2.0 [−4.8, 0.7] | $2.4 \times 10^{-1}$ |
| −1→1 | Control vs EphA3-Fc + KYL | 20,2 | 21,8 | −1.7 [−4.5, 1.1] | $3.4 \times 10^{-1}$ |
| −1→1 | Control vs βMCD | 20,2 | 19,1 | 1.1 [−1.6, 3.7] | $5.3 \times 10^{-1}$ |
| −1→1 | Control vs EphA3-Fc + βMCD | 20,2 | 21,1 | −1.0 [−3.7, 1.7] | $5.3 \times 10^{-1}$ |
| −1→1 | EphA3-Fc + βMCD vs EphA3-Fc + KYL | 21,1 | 21,8 | −0.7 [−3.6, 2.2] | $6.3 \times 10^{-1}$ |
| 1→−1 | EphA3-Fc vs EphA3-Fc + KYL | 13,7 | 21,6 | −7.9 [−10.7, −5.1] | $6 \times 10^{-7}$ |
| 1→−1 | Control vs EphA3-Fc | 21,0 | 13,7 | 7.4 [4.7, 10.0] | $6.2 \times 10^{-7}$ |
| 1→−1 | EphA3-Fc vs EphA3-Fc + βMCD | 13,7 | 20,9 | −7.3 [−10.0, −4.6] | $8.7 \times 10^{-7}$ |
| 1→−1 | EphA3-Fc vs βMCD | 13,7 | 19,6 | −6.0 [−8.6, −3.3] | $4.6 \times 10^{-5}$ |
| 1→−1 | βMCD vs EphA3-Fc + KYL | 19,6 | 21,6 | −1.9 [−4.8, 0.9] | $3.7 \times 10^{-1}$ |
| 1→−1 | Control vs βMCD | 21,0 | 19,6 | 1.4 [−1.3, 4.1] | $5.1 \times 10^{-1}$ |
| 1→−1 | βMCD vs EphA3-Fc + βMCD | 19,6 | 20,9 | −1.3 [−4.1, 1.4] | $5.1 \times 10^{-1}$ |
| 1→−1 | EphA3-Fc + βMCD vs EphA3-Fc + KYL | 20,9 | 21,6 | −0.6 [−3.5, 2.3] | $7.9 \times 10^{-1}$ |
| 1→−1 | Control vs EphA3-Fc + KYL | 21,0 | 21,6 | −0.5 [−3.4, 2.3] | $7.9 \times 10^{-1}$ |
| 1→−1 | Control vs EphA3-Fc + βMCD | 21,0 | 20,9 | 0.1 [−2.6, 2.8] | $9.5 \times 10^{-1}$ |
| 1→0 | βMCD vs EphA3-Fc + KYL | 5,9 | 3,0 | 2.8 [1.4, 4.3] | $1.6 \times 10^{-3}$ |
| 1→0 | EphA3-Fc vs βMCD | 3,3 | 5,9 | −2.5 [−4.0, −1.0] | $7.3 \times 10^{-3}$ |
| 1→0 | Control vs βMCD | 4,1 | 5,9 | −1.8 [−3.2, −0.3] | $5.5 \times 10^{-2}$ |
| 1→0 | EphA3-Fc + βMCD vs EphA3-Fc + KYL | 4,5 | 3,0 | 1.5 [0.2, 2.9] | $6.9 \times 10^{-2}$ |
| 1→0 | βMCD vs EphA3-Fc + βMCD | 5,9 | 4,5 | 1.3 [−0.2, 2.8] | $1.6 \times 10^{-1}$ |
| 1→0 | EphA3-Fc vs EphA3-Fc + βMCD | 3,3 | 4,5 | −1.2 [−2.6, 0.2] | $1.6 \times 10^{-1}$ |
| 1→0 | Control vs EphA3-Fc + KYL | 4,1 | 3,0 | 1.1 [−0.2, 2.3] | $1.6 \times 10^{-1}$ |
| 1→0 | Control vs EphA3-Fc | 4,1 | 3,3 | 0.7 [−0.6, 2.1] | $3.6 \times 10^{-1}$ |
| 1→0 | Control vs EphA3-Fc + βMCD | 4,1 | 4,5 | −0.5 [−1.8, 0.9] | $5.6 \times 10^{-1}$ |
| 1→0 | EphA3-Fc vs EphA3-Fc + KYL | 3,3 | 3,0 | 0.3 [−1.0, 1.6] | $6.3 \times 10^{-1}$ |
| 0→1 | βMCD vs EphA3-Fc + KYL | 6,2 | 2,5 | 3.7 [2.3, 5.1] | $6.7 \times 10^{-6}$ |
| 0→1 | EphA3-Fc vs βMCD | 2,9 | 6,2 | −3.3 [−4.8, −1.8] | $1.5 \times 10^{-4}$ |
| 0→1 | βMCD vs EphA3-Fc + βMCD | 6,2 | 4,2 | 2.0 [0.5, 3.5] | $2.1 \times 10^{-2}$ |
| 0→1 | Control vs EphA3-Fc + KYL | 4,4 | 2,5 | 1.9 [0.6, 3.1] | $1.3 \times 10^{-2}$ |
| 0→1 | Control vs βMCD | 4,4 | 6,2 | −1.8 [−3.3, −0.3] | $3 \times 10^{-2}$ |
| 0→1 | EphA3-Fc + βMCD vs EphA3-Fc + KYL | 4,2 | 2,5 | 1.7 [0.5, 3.0] | $2.1 \times 10^{-2}$ |
| 0→1 | Control vs EphA3-Fc | 4,4 | 2,9 | 1.5 [0.2, 2.8] | $4.7 \times 10^{-2}$ |
| 0→1 | EphA3-Fc vs EphA3-Fc + βMCD | 2,9 | 4,2 | −1.3 [−2.6, 0.0] | $7.7 \times 10^{-2}$ |
| 0→1 | EphA3-Fc vs EphA3-Fc + KYL | 2,9 | 2,5 | 0.4 [−0.8, 1.6] | $5.5 \times 10^{-1}$ |
| 0→1 | Control vs EphA3-Fc + βMCD | 4,4 | 4,2 | 0.2 [−1.2, 1.5] | $8 \times 10^{-1}$ |

**Supplementary Table 1. Complete pairwise comparisons of stability and transition frequencies following EphA3-Fc, βMCD, and KYL treatments.** All pairwise comparisons are reported, including both statistically significant and non-significant results, for overall stability and for each transition category (1→1, −1→−1, −1→1, 1→−1, 1→0, and 0→1). Group 1 (%) and

Group 2 (%) indicate the percentage of observations assigned to the corresponding outcome or transition category in each experimental group. Differences are expressed in percentage points (Δ p.p.; Group 1 − Group 2), together with their 95% confidence intervals (95% CI). q values were adjusted for multiple comparisons using a false discovery rate (FDR) correction. An FDR-adjusted q value $< 0.05$ was considered statistically significant.

**SUPPLEMENTARY RESULTS: Exploratory relationships among guidance and growth parameters**

**Exploratory relationships among guidance and growth parameters**

To integrate axonal growth dynamics and guidance, we performed exploratory analyses to determine whether the terminal deviation angle (β) or average growth velocity was associated with variables describing the initial state of the axon or its spatial context during the assay. The variables examined included initial axon length, initial orientation relative to the stimulus, relative position within the molecular gradient, average growth velocity, dynamic persistence, and pause fraction. These variables were considered descriptors of the axonal state and experimental context at the time of acquisition, rather than intrinsic determinants of the guidance response. The analyses were intended to identify potential sources of heterogeneity or bias and were not designed as formal tests of mediation or causality. We first examined whether initial orientation and relative gradient position were associated with average growth velocity. We then assessed whether the deviation angle (β) was associated with initial orientation, initial length, relative gradient position, average velocity, dynamic persistence, or pause fraction.

### Initial Angle and Average Velocity

To assess the influence of the initial axonal angle relative to the molecular gradient on average growth velocity, axons were classified based on whether their initial direction was "toward" or "away from" the gradient. Average axonal growth velocity did not differ between axons oriented "toward" vs. "away from" the molecular gradients in nasal RGCs under any tested condition—including control, EphA3-Fc, EphA3-Fc + KYL, βMCD, and EphA3-Fc + βMCD (all comparisons were non-significant, *n.s.*; Supplementary Figure 4A, 4B). These findings indicate that the gradients and modulators evaluated do not introduce a kinetic bias dependent on the direction of axonal incidence.

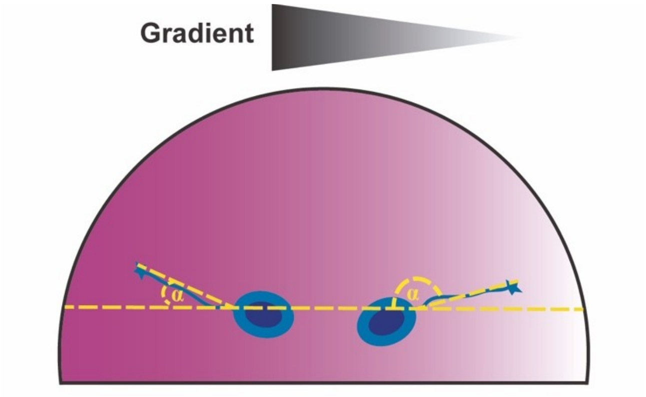


**Supplementary Figure 4A. Determination of the initial angle (α).** RGCs (blue circles) are positioned on a semicircle within a lateral concentration gradient (dark → light). The angle α is measured between the basal reference line (gradient direction) and the initial axonal trajectory (yellow dashed lines). Axons with angles from 0° to 90° were designated as "toward" the gradient, whereas those with angles >90° to 180° were designated as "away from" the gradient to correlate initial angle with growth velocity.

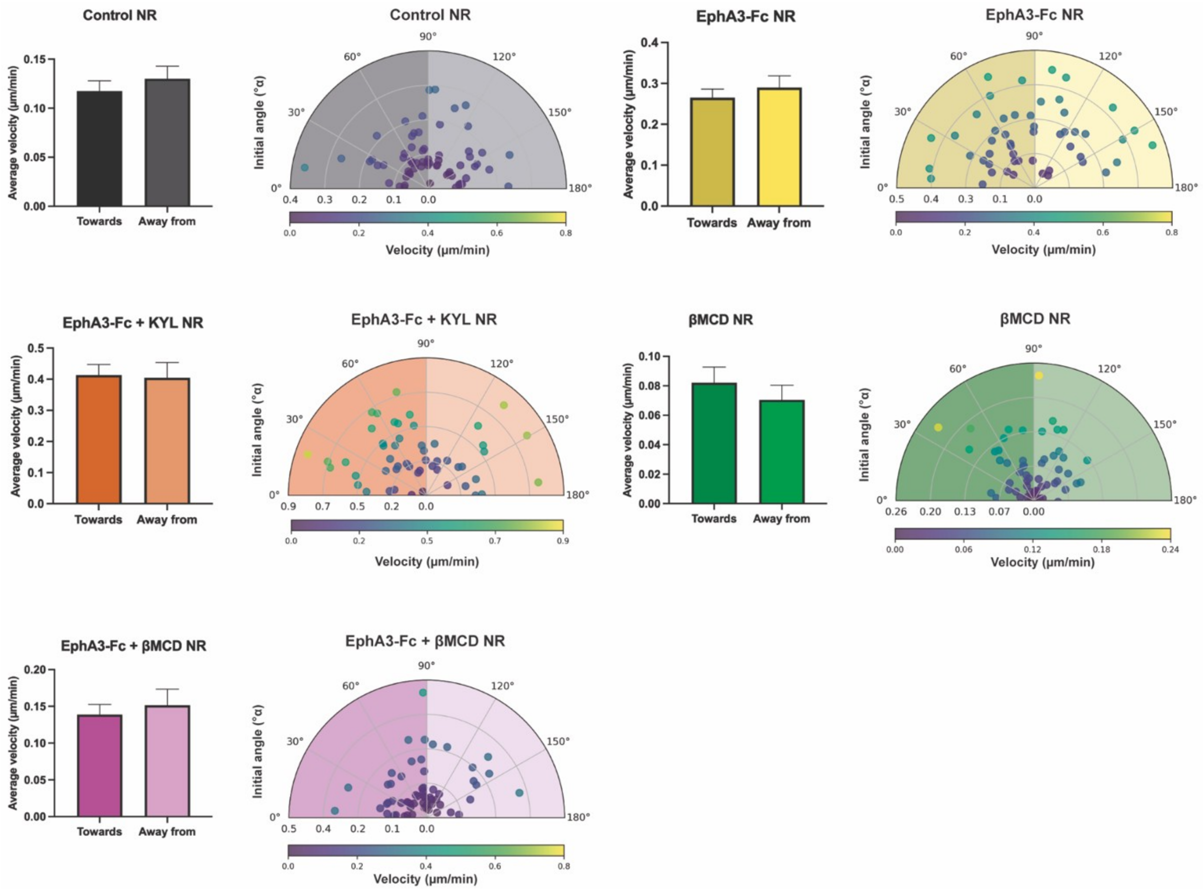


**Supplementary Figure 4B. Axonal growth velocity as a function of gradient incidence and treatment.** For each condition, the following are shown: Bar graphs (left of each pair): mean velocity ± SEM of axons advancing toward (0°–90°) versus away from (>90°–180°) the gradient (Mann-Whitney U test; "ns" = non-significant). To the right, a semicircular polar plot represents each individual axon: the radial angle corresponds to the initial direction (0° = toward the gradient, 180° = away from), the distance from the origin reflects velocity magnitude, and the color scale (bottom bar of each plot) encodes that same velocity in µm $min^{-1}$. Background circles outline radii equivalent to 25%, 50%, and 75% of the maximum velocity observed in the corresponding condition.

### Initial Angle and Deviation Angle

To determine whether the initial axonal orientation (initial angle α) relative to the molecular gradient influences terminal targeting (deviation angle β), the correlation between the initial growth angle (°) and the deviation angle (°) was analyzed in nasal and temporal RGCs across all evaluated conditions (control, EphA3-Fc, βMCD, EphA3-Fc + βMCD, EphA3-Fc + KYL). No association was observed between the initial angle (α) and deviation angle (β): regression slopes were near zero, with 95% confidence intervals (95% CI) including the null value, and Spearman correlations were non-significant ($p > 0.05$) (Supplementary Figure 5). The dispersion of β was comparable across the entire range of α, showing no consistent upward or downward trends. Taken together, these results indicate that the angular deviation (deviation angle β) exhibited by axons under the influence of different molecular gradients does not depend on their starting orientation (initial angle α) at the time of measurement.

From a guidance perspective, this suggests that, within the tested timeframes and conditions, the growth cone can reorient in response to guidance cues independently of its initial heading. Thus, gradient information appears to operate by modulating turning probability rather than imposing a deviation rate proportional to the starting angle.

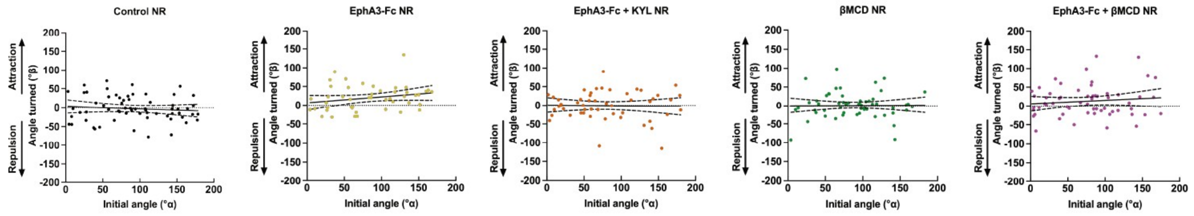


**Supplementary Figure 5. Lack of correlation between initial growth angle (α) and deviation angle (β) for nasal and temporal RGCs across all treatments, including control, EphA3-Fc, βMCD, EphA3-Fc + βMCD and EphA3-Fc + KYL.** Statistical analysis (Spearman), covering attraction (positive values) and repulsion (negative values), indicating the regression slopes did not differ from zero, demonstrating independent axonal guidance behavior ($p > 0.05$). Each point represents an individual axon; the solid line indicates the linear regression and the discontinuous lines represent the 95% CI.

### Gradient Concentration Zone and Axonal growth velocity.

To determine whether the relative molecular concentration within the gradient surrounding the analyzed neurons influences axonal growth velocity, growth rates were compared among RGCs located across three molecular gradient zones (high, medium, and low concentration) (Fig. 6). No significant differences in average axonal growth velocity were observed among the different gradient zones for any condition (Supplementary Figure 6). These results demonstrate that the types of gradients utilized (with their respective slopes) do not establish significant differences in axonal growth velocity among neurons located within the three analyzed regions. These findings provide no evidence that neuron distribution across gradient zones accounts for the velocity differences observed in other analyses; however, subtle effects cannot be excluded with the current design.

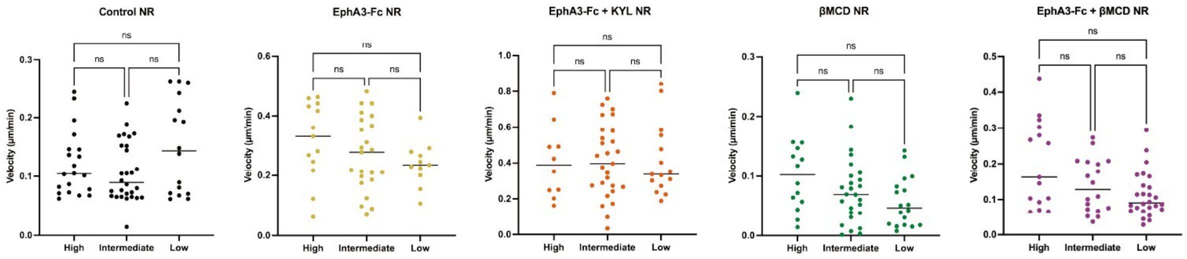


**Supplementary Figure 6. Average axonal growth velocity as a function of gradient concentration.** Average axonal velocity (µm/min) was evaluated based on the gradient zone where the axons were located (high, medium, or low relative concentration). Axons from nasal (NR) RGCs were analyzed under control conditions or in the presence of EphA3-Fc, βMCD, EphA3-Fc + βMCD, and EphA3-Fc + KYL. Data are shown as group scatter plots with ± SEM; the horizontal line indicates the median. Statistical comparisons were performed using a non-parametric Kruskal–Wallis ANOVA followed by Dunn's post hoc test. No significant differences were detected in any case (ns, $p > 0.05$).

### Gradient Concentration Zone and Deviation Angle

To determine whether the relative molecular concentration within the gradient surrounding the analyzed neurons influences terminal axonal targeting, deviation angles (β) were compared among RGCs located across three molecular gradient zones (high, medium, and low concentration) (Supplementary Figure 7). No significant differences in β angles were detected among the high, medium, or low concentration zones under any condition. Within the utilized gradient type, the deviation angle did not depend on the relative position of the axon within the gradient, ruling out potential bias based on the proportion of neurons analyzed in each region.

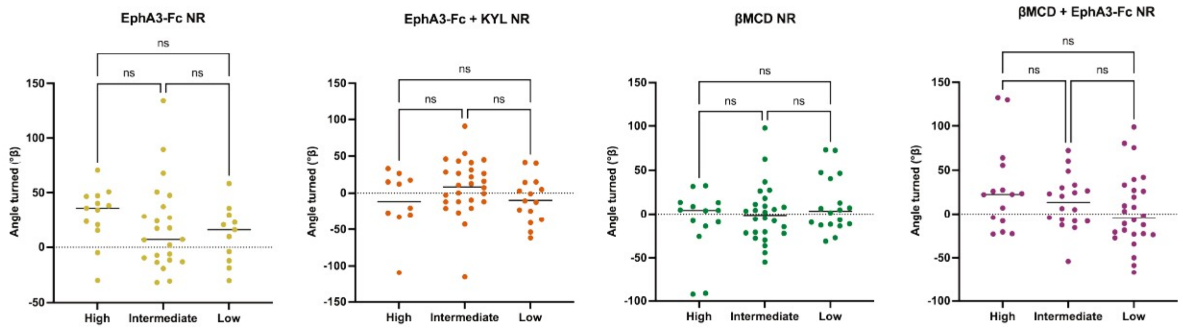

**Supplementary Figure 7. Deviation Angle as a function of gradient concentration.** Axon deviation angles for nasal retina (NR) were measured across high, medium, and low gradient concentrations under control or experimental conditions (EphA3-Fc, EphA3-Fc + KYL, βMCD, and βMCD + EphA3-Fc). Statistical comparisons were performed using a non-parametric Kruskal–Wallis ANOVA followed by Dunn's post hoc test. The analysis, covering attraction (positive values) and repulsion (negative values), indicates no significant directional changes based on axon position within the gradient (ns, $p > 0.05$).

### Initial axonal length and deviation angle

In order to evaluate whether the state of axonal length influences the level of axonal growth velocity, analyses of deviation angle β relative to initial axonal length were performed. They reveal no statistical association across experimental conditions, indicating the degree of angular deviation is independent of the initial axon length (Supplementary Figure 8). The findings suggest that guidance cue sensitivity and resulting axon steering are driven by molecular mechanisms independent from the morphological state of elongation at the time of detection. This independence enhances the robustness of topographic mapping.

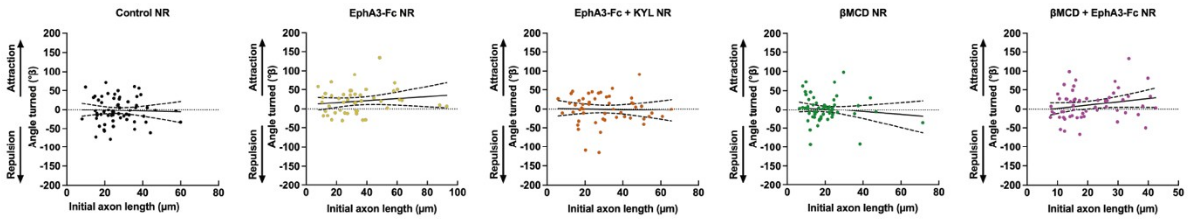

**Supplementary Figure 8. Relationship between initial axon length and angular deviation.** The correlation between initial axon length (μm) and deviation angle (°) was analyzed in nasal RGCs (NR) under control conditions or stimulated with EphA3-Fc, EphA3-Fc + KYL, βMCD, and βMCD + EphA3-Fc. In all treatments, the regression slope did not differ significantly from zero, and correlation coefficients were non-significant (Spearman, $p > 0.05$), indicating the absence of a linear relationship between both parameters.

### Initial axonal length and axonal growth velocity.

In order to evaluate whether the axonal length conditions the axonal growth velocity, the existence of correlation between these parameters was analyzed (Supplementary Figure 9). Only nasal RGC with EphA3-Fc + KYL showed a significant negative correlation between initial axon length and average velocity (ρ = −0.53, p < 0.0001).

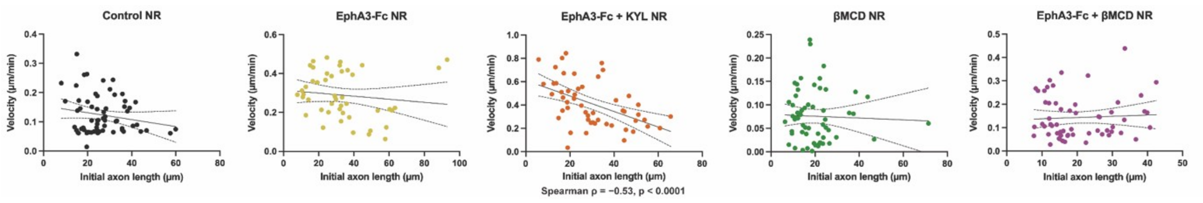


**Supplementary Figure 9. Relationship between initial axon length and axonal growth velocity.** The correlation between initial axon length (μm) and axonal growth velocity was analyzed in nasal RGCs (NR) under control conditions or stimulated with EphA3-Fc, EphA3-Fc + KYL, βMCD, EphA3-Fc + βMCD. Spearman's rank correlation coefficient was calculated for each condition. Only EphA3-Fc + KYL NR showed a significant negative correlation between initial axon length and average velocity (ρ = −0.53, p < 0.0001).

### Axonal growth velocity and deviation Angle

To evaluate whether growth kinetics condition turning capacity, the association between elongation velocity and deviation angle (β) was analyzed (Supplementary Figure 10). Across all conditions (control, EphA3-Fc, EphA3-Fc + KYL, βMCD, EphA3-Fc+ βMCD; NR: Nasal Retina), regression slopes were near zero and correlations were non-significant. Taken together, no evidence was found that turning magnitude (β) depends on how fast the axon grows within the evaluated timeframes and experimental ranges. The absence of robust correlations between velocity and β suggests a functional decoupling between axonal orientation and propulsion.

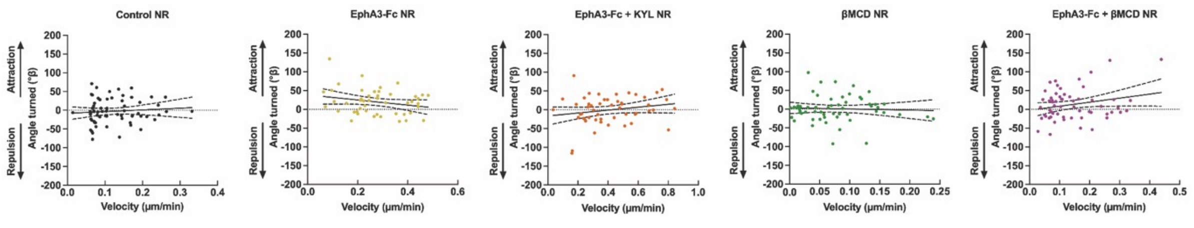


**Supplementary Figure 10. Deviation angle as a function of average axonal velocity.** The correlation between average elongation velocity (μm/min) and deviation angle (°) was evaluated in nasal RGCs (NR) under control conditions or in the presence of EphA3-Fc, EphA3-Fc + KYL, βMCD, EphA3-Fc + βMCD. In no condition did the slope differ from zero, nor did the correlation coefficient reach statistical significance (*p* > 0.05).

### Dynamic persistence and deviation angle

In order to evaluate whether the level of advances and retraction intervals influences the terminal guidance, we correlated Dynamic Persistence Index (DPI) with the deviation angle (β).

The Dynamic Persistence Index (DPI) was calculated as the ratio of net axonal advances to total movement events (DPI= *N* advances – *N* retractions / *N* advances + *N* retractions), where "N advances" is the number of intervals in which the axon advance (>0) and "N retractions" is the number of intervals in which the axon retracts (<0). Pauses (0) were not included. DPI value is between -1 and +1 (-1: only retractions, 0: balanced, +1: only advances). DPI is independent from timelapse duration and from mean axon growth velocity.

DPI showed no significant ($p$ > 0.05) correlation with terminal deviation angle (β) across experimental conditions (Supplementary Figure 11). Regardless of high or low persistence, axon growth displayed varied responses (attraction, repulsion, or no deviation), indicating that deviation magnitude does not depend on dynamic persistence within the evaluated ranges.

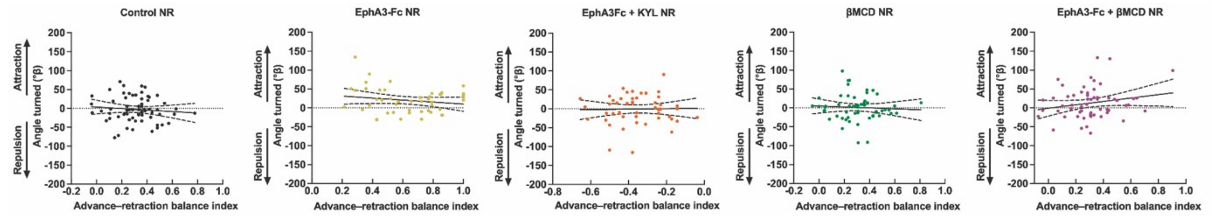


**Supplementary Figure 11. Relationship between the dynamic persistence of axonal growth and the deviation angle.** The plot includes nasal RGC (NR) controls, as well as conditions with EphA3-Fc, EphA3-Fc + KYL, βMCD, EphA3-Fc + βMCD. In dynamic persistence (advances − retractions)/(advances + retractions), 1 indicates only advances and 0 depicts equal number of advances and retractions. Each graphic shows the individual values of persistence (X axis) against deviation angle at 2 h of time-lapse (Y axis): positive = attraction towards the gradient, negative = repulsion. In no condition did the slope differ from zero, nor did the correlation coefficient reach statistical significance ($p$ > 0.05).

### Pause Fraction and Deviation Angle

To evaluate whether growth cone immobility influences terminal guidance, we correlated the pause fraction with the deviation angle (β) under conditions where pausing behaviors exhibited significant differences (Supplementary Figure 12). We define the pause fraction as follows:

Pause Fraction =$N$ pauses / $N$ advances + $N$ retractions + $N$ pauses

The pause fraction is an index that quantifies the proportion of time (or number of events) an axon remains immobile relative to its total motility behaviors during the time-lapse recording. By definition, its value ranges from 0 to 1 (0: never pauses; 1: remains stationary throughout the entire recording).

Across all analyzed conditions, the slopes approached zero, and the correlations were not statistically significant (p > 0.05). The magnitude of (β) remained highly variable across the entire range of the pause fraction, showing no systematic trends. Thus, we found no evidence that the magnitude of the deviation (β) depends on the relative pausing time within the experimental windows and ranges evaluated.

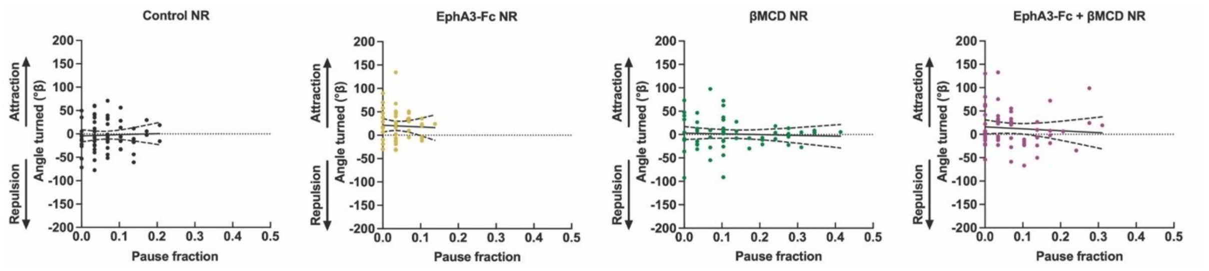


**Supplementary Figure 12. Relationship between pause fraction and axonal directional deviation.** The pause fraction was defined as pauses/(advances + retractions + pauses); values near 0 indicate axons that are active nearly the entire time, whereas high values reflect extended periods of immobility. The X-axis was truncated at 0.5 because no conditions yielded fractions above this threshold. For each treatment (EphA3-Fc, βMCD, EphA3-Fc + βMCD) and control, the pause fraction was plotted against the deviation angle at 2 h (Y-axis; positive values indicate attraction toward the gradient, and negative values indicate repulsion). The solid line represents the simple linear regression, and the dashed lines indicate the 95% confidence interval. In all cases, both the slope and the correlation coefficient were not statistically significant ($p > 0.05$).

Overall, deviation angle (β) showed no consistent association with initial orientation, initial axon length, relative position within the gradient, average growth velocity, dynamic persistence, or pause fraction within the conditions and ranges examined. Initial orientation and gradient position were likewise not associated with average velocity. A significant negative association between initial axon length and average velocity was detected specifically in nasal RGCs treated with EphA3-Fc + KYL; this isolated result should be interpreted cautiously in the context of the multiple exploratory comparisons performed. These findings suggest that the measured baseline and growth-dynamic variables are unlikely to account systematically for the observed guidance responses, while not excluding smaller effects or associations restricted to individual experimental conditions.
